# The Arthropoda-specific Tramtrack group BTB protein domains use previously unknown interface to form hexamers

**DOI:** 10.1101/2022.09.01.506177

**Authors:** Artem N. Bonchuk, Konstantin I. Balagurov, Rozbeh Baradaran, Konstantin M. Boyko, Nikolai N. Sluchanko, Anastasia M. Khrustaleva, Anna D. Burtseva, Olga V. Arkova, Karina K. Khalisova, Vladimir O. Popov, Andreas Naschberger, Pavel G. Georgiev

## Abstract

BTB (Bric-a-brack, Tramtrack and Broad Complex) is a diverse group of protein-protein interaction domains found within metazoan proteins. Transcription factors contain a dimerizing BTB subtype with a characteristic N-terminal extension. The Tramtrack group (TTK) is a distinct type of BTB domain, which can multimerize. Single-particle cryo-EM microscopy revealed that the TTK-type BTB domains assemble into a hexameric structure consisting of three canonical BTB dimers connected through a previously uncharacterized interface. We demonstrated that the TTK-type BTB domains are found only in Arthropods and have undergone lineage-specific expansion in modern insects. The *Drosophila* genome encodes 24 transcription factors with TTK-type BTB domains, whereas only four have non-TTK-type BTB domains. Yeast two-hybrid analysis revealed that the TTK-type BTB domains have an unusually broad potential for heteromeric associations presumably through dimer-dimer interaction interface. Thus, the TTK-type BTB domains are a structurally and functionally distinct group of protein domains specific to Arthropodan transcription factors.

## Introduction

BTB, also known as POZ (Pox-virus and zinc-finger), is an evolutionarily conserved domain that was originally found in the *Drosophila* proteins bric-a-brac, Tramtrack and broad complex (Zollman et al., 1994). BTB proteins have been identified in poxviruses and eukaryotes, and have various functions including regulation of transcription, chromatin remodeling, cytoskeletal function, ion transport, and ubiquitination/degradation of proteins (Perez-Torrado et al., 2006; Stogios et al., 2005). BTB domains are 100–120 aa in size and the core structure consists of five α-helices and three β-strands (Ahmad et al., 1998; Bonchuk et al., 2023; Li et al., 1999). In addition to this core, different subclasses of BTB proteins include N- and C-terminal BTB extension regions that facilitate protein-specific functions. As a result, the BTB fold is a versatile scaffold that participates in a variety of family-specific protein-protein interactions. In metazoans, BTB domains of transcription factors contain an amino-terminal extension that enables homodimerization (Ahmad et al., 1998; Ahmad et al., 2003; Bonchuk et al., 2023). Most members of this family contain C2H2-type zinc fingers (BTB-C2H2). In humans, 156 genes are predicted to encode BTB domain-containing proteins, of which 49 possess between 2 and 14 C2H2 domains (Siggs and Beutler, 2012). BTB-C2H2 proteins act as classical transcription factors, binding to chromatin and participating in the regulation of transcription. In insects, some transcription factors contain BTB domains in combination with other DNA-binding domains such as Pipsqueak (PSQ, BTB PSQ) (Siegmund and Lehmann, 2002) and helix-turn-helix (HTH) . Several BTB-containing transcription factors contain poorly characterized FLYWCH (named after characteristic sequence motif) domains that may be involved in the interaction with either DNA, RNA, or proteins (Beaster-Jones and Okkema, 2004; Melnikova et al., 2017).

In mammals, BTB-C2H2 transcription factors are required for the development of lymphocytes, fertility, skeletal morphogenesis, and neurological development (Chaharbakhshi and Jemc, 2016). The well-characterized BTB domains form tightly intertwined dimers and possess a peptide-binding groove, which is responsible for the interaction with various transcription factors and co-repressor complexes (Ahmad et al., 2003; Ghetu et al., 2008; Vogelmann et al., 2014; Zacharchenko and Wright, 2021). In most cases, mammalian C2H2 transcription factors have BTB domains which exclusively form homodimers. An exception to this rule is the C2H2 protein Miz-1, whose BTB domain forms tetramers (Stead et al., 2007). Several BTB domains are capable of heterodimer formation (Olivieri et al., 2021). The structural basis for heterodimerization has been studied using chimeric Bcl6/Miz-1 and Miz-1/NAC1 assembly (Stead and Wright, 2014); however, there is evidence that formation of such heterodimers *in vivo* is prevented by co-translational dimer assembly and the quality-control protein degradation machinery (Bertolini et al., 2021; Mena et al., 2020; Mena et al., 2018).

In *Drosophila*, a number of C2H2 proteins with a BTB domain have been described, some containing a typical BTB domain, such as CP190 and CG6792 (Maeng et al., 2012). The most well-studied of these, CP190, has four C2H2 zinc finger domains that appear to be involved in protein-protein interactions rather than DNA binding (Oliver et al., 2010). The N-terminal BTB/POZ domain of CP190 forms stable homodimers (Bonchuk et al., 2011; Oliver et al., 2010; Plevock et al., 2015; Sabirov et al., 2021b; Vogelmann et al., 2014). CP190 is required for the activity of housekeeping promoters and insulators (Bartkuhn et al., 2009; Sabirov et al., 2021a). CG6792 is similar to mammalian BTB-C2H2 proteins, has seven DNA-binding zinc fingers, and is involved in wing development (Maeng et al., 2012).

Most of the well-characterized *Drosophila* BTB transcription factors, including GAF, Mod(mdg4), LOLA, Broad-complex (BR-C), Batman, Pipsqueak, and Bric-a-brac (Bab), have TTK-type BTB domains (Bonchuk et al., 2011; Zollman et al., 1994). Proteins from this group often have important functions in transcription regulation, development, and chromosome architecture (Chaharbakhshi and Jemc, 2016). Many TTK-group proteins, such as BR-C, TTK and Bab, are critical regulators of development that function as transcriptional repressors (Bradley and Andrew, 2001; Chaharbakhshi and Jemc, 2016; Mukai et al., 2007; Silva et al., 2016). Several have been implicated in chromatin architectural function, acting as a component of a chromatin insulator complex (Mod(mdg4)) or recruiting chromatin remodeling complexes (GAF, Pipsqueak) (Huang et al., 2002; Lomaev et al., 2017). Recently, it was shown that GAF is a pioneer factor that acts as a stable mitotic bookmarker during zygotic genome activation during *Drosophila* embryogenesis (Bellec et al., 2022; Tang et al., 2022).

TTK-type BTB domains contain a highly conserved N-terminal FxLRWN motif, where x is a hydrophilic residue (Bonchuk et al., 2011). Several TTK-type BTB domains can selectively interact with each other and form multimers (Bonchuk et al., 2011). Although there are more than 30 crystal structures of non-TTK BTB domains that form stable homodimers (Ahmad et al., 1998; Stogios et al., 2007; Stogios et al., 2010; Stogios et al., 2005; Vogelmann et al., 2014), the structures of the TTK-type BTB domains and the structural basis for their multimerization remain unknown.

In this study, we investigated the presence of TTK-type and canonical BTB domains in transcription factors from various phylogenetic groups of Bilateria. The TTK-type BTB domains were found only in Arthropodan transcription factors and underwent lineage-specific expansion in insects. Using an integrative structural biology approach, we built a structural model of the TTK-group BTB hexameric assembly, validating it using MALS, SAXS, cryo-EM microscopy, and site-directed mutagenesis. Finally, we found an unusual potential for these domains to form heteromultimers, which likely involves a dimer-dimer interaction interface.

## Results

### Most BTB domains of transcription factors in *Drosophila melanogaster* are of TTK-type

The *Drosophila melanogaster* genome contains 28 genes encoding transcription factors with BTB domains (Figure 1). Many genes encode several BTB-containing protein isoforms, which differ in their C-terminal sequences. At two exceptional loci, *mod(mdg4)* and *lola*, multiple isoforms are formed by *trans*-splicing, the mechanism of which is still unclear (Tikhonov et al., 2018). The *mod(mdg4)* locus encodes at least 30 isoforms, most of which have FLYWCH domains (Buchner et al., 2000). Seventeen of the 20 isoforms produced by the *lola* locus contain different C-terminal C2H2 zinc-fingers (Horiuchi et al., 2003).

**Figure 1.**
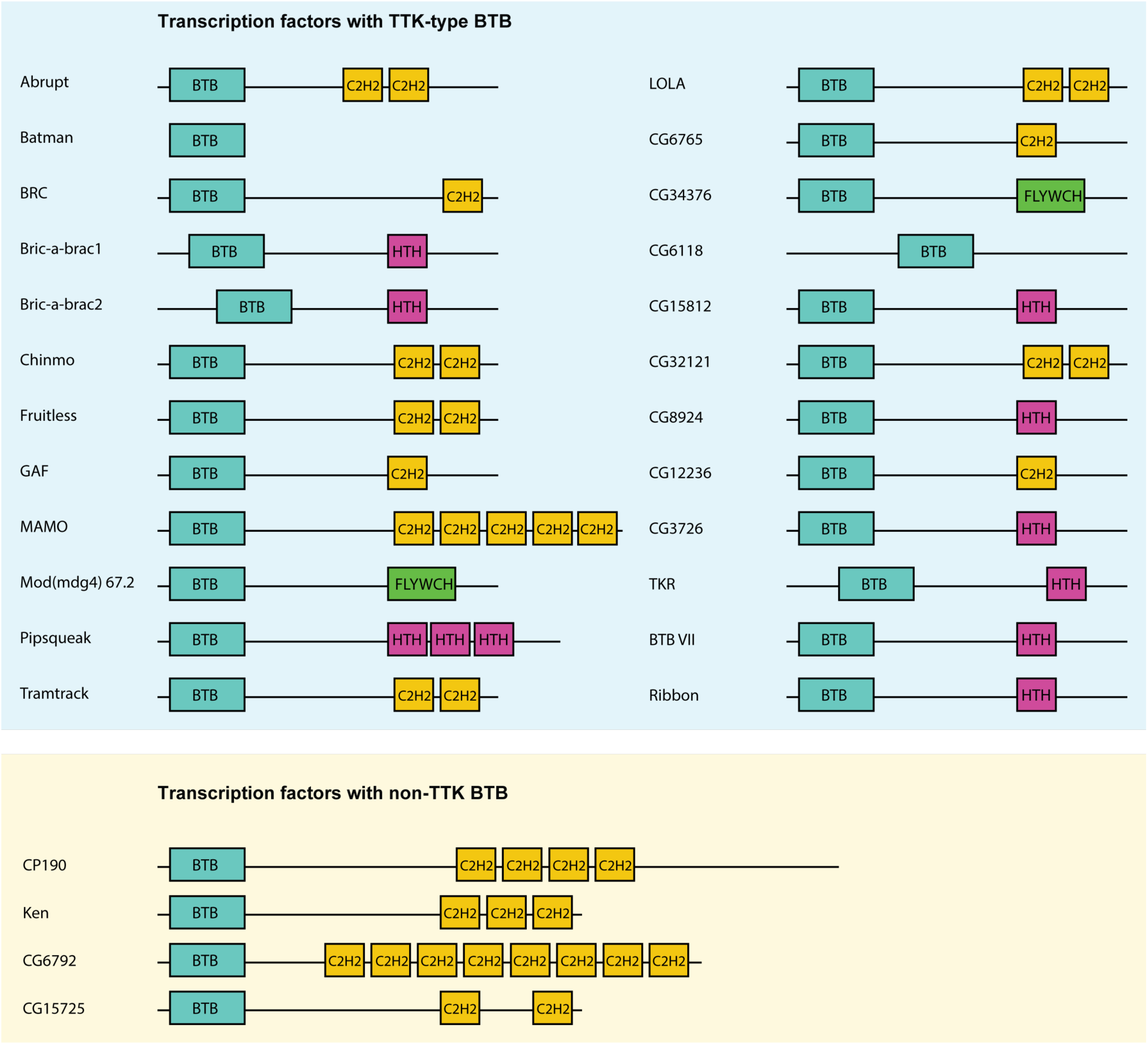
Schematic representation of Drosophila transcription factors with BTB domains. Only one representative isoform is shown for each gene.

A characteristic feature of the TTK-type BTB domain is the presence of the FxLRWN motif at the N-terminus. Comparison of the amino acid sequence of BTBs from 28 *Drosophila* BTB-containing transcription factors showed that 24 BTB domains contain the characteristic “TTK motif” (Figure 2a and 2b). The TTK-type BTB domains are usually located at N-termini of proteins, however, in two proteins (CG6118, bric-a-brac2), the BTB domains are located in the middle of the protein (Figure 1). As an exception, Batman consists of only the BTB domain. Four transcription factors (CP190, CG6792, CG15725 and Ken) have BTB domains without a TTK motif (Figure 1).

**Figure 2.**
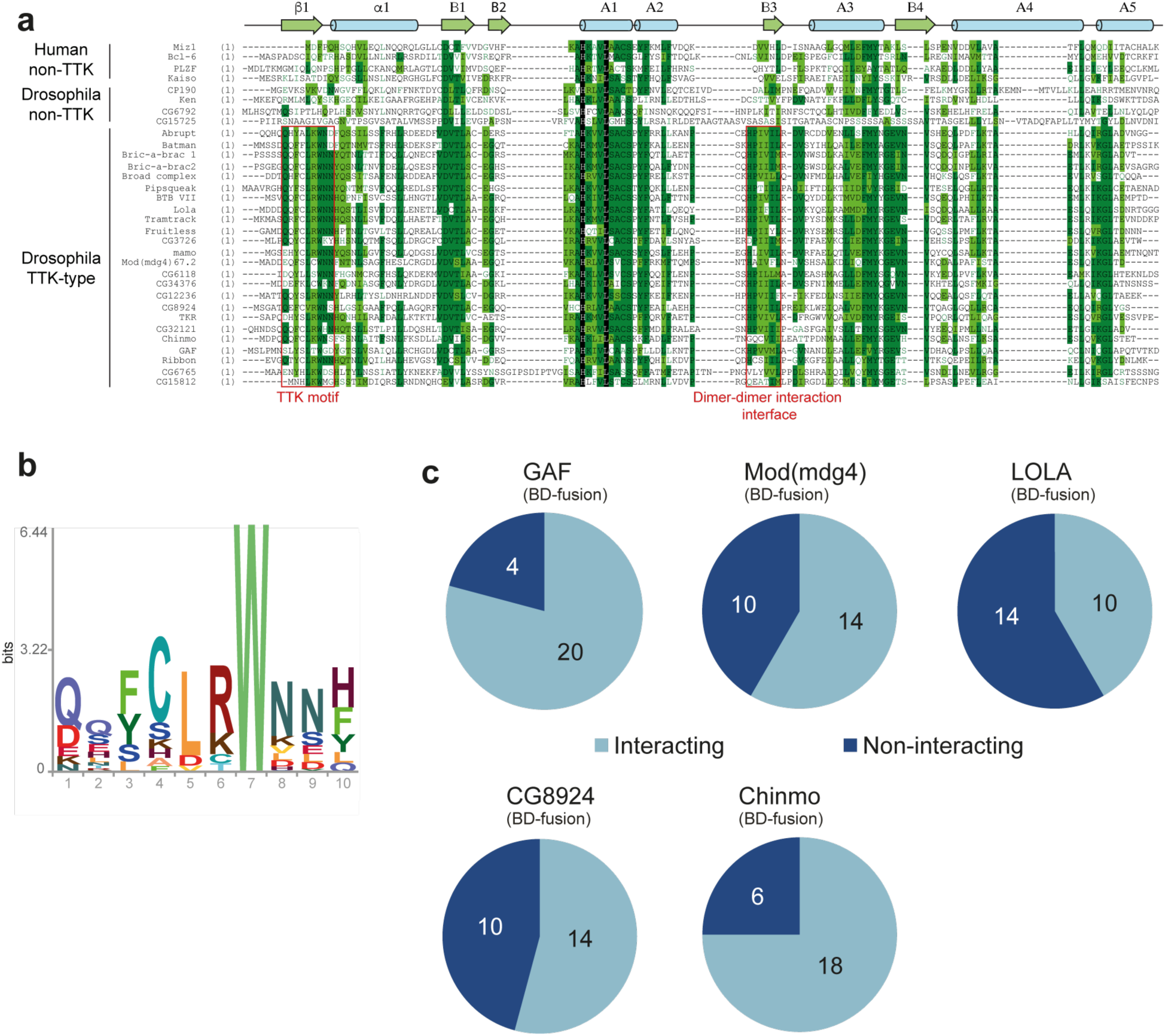
Characterization of the TTK-type BTB domains. **a** Multiple sequence alignment of BTB domains from *Drosophila* transcription factors and a few human BTB domains with known 3D structures. Secondary structure elements are labeled according to (Stogios et al., 2005). **b** HMM-profile model for the TTK motif obtained for 14 Diptera species. **c** Testing of the GAF, Mod(mdg4), and LOLA BTB domains for interaction with all TTK-type BTB domains found in *Drosophila melanogaster*. Original data are shown in Supplementary Table S3.

### The TTK-type BTB domains display a broad heteromeric interaction propensity

The characteristic feature of the TTK-type BTB domains is their ability to selectively interact with each other (Bonchuk et al., 2011). We tested interaction of all TTK-type BTB domains with BTB domains from Mod(mdg4), LOLA, Chinmo, CG8924 and GAF in a yeast two-hybrid assay (Y2H; Figure 2c, Supplementary Table S3 and Supplementary Figure S2). We found that all BTB domains can interact with themselves. 23 out of 24 TTK-type BTB domains also interacted with at least one of GAF, LOLA, or Mod(mdg4) BTB domains. GAF, Mod(mdg4), LOLA and CG8924 all interacted with at least 10 of the 24 TTK-type BTB domains, while Chinmo interacted with 18 TTK-type BTB domains (Figure 2c, Supplementary Table S3). The homology between interacting domains was mostly in the range 35–47%, but could be as low as 25%. No obvious relationship between homology and heteromeric interaction ability was found (Supplementary Table S4).

We also tested interaction between four non-TTK BTB domains (CP190, Ken, CG15275 and CG6792 (dPLZF)) and did not find heteromeric interaction between them (Supplementary Table S5). The BTBs of CP190 and CG6792 formed homodimers *in vitro,* like all other classical BTB domains (Supplementary Figure S3). The Ken and CG15725 BTB domains were insoluble after bacterial expression. Using Y2H, we detected only a small number of interactions between the non-TTK BTB domains of CP190 and Ken with the TTK-type BTB domains: CP190 interacted with GAF and Ribbon, while Ken interacted with Batman and Mamo (Supplementary Table S6 and Supplementary Figure S4). These results confirm that the TTK-type domains are functionally distinct from classical BTB domains.

### Cryo-electron microscopy reveals hexameric assembly of the BTB domain of CG6765 protein through a previously uncharacterized interface

To test the ability of TTK-type BTB domains to multimerize and shed light on their possible structure, we screened all TTK-type BTB domains from *Drosophila melanogaster* for possibility of high-yield expression and soluble purification for subsequent structural analysis.

We expressed all 24 TTK-type *Drosophila* BTB domains as TEV-cleavable thioredoxin fusions. Out of 24 BTBs, only ten domains were soluble and stable after TEV-cleavage. Eight BTB domains appeared as a single peak on size-exclusion chromatography (SEC) with apparent Mw 90–150 kDa, Batman BTB eluted as a dimer (Supplementary Figure S5), while CG32121 BTB formed multiple peaks. Unfortunately, most of these BTBs had low solubility (∼1 mg/mL) and, therefore, were not suitable for structural studies.

Only three BTB domains (CG6765^1-133^, CG32121^1-147^ and LOLA^1-120^) were soluble at concentrations above 5 mg/mL. CG32121^1-147^ was excluded from further analysis due to its heterogeneous oligomeric state. Multiple crystallization trials led to nicely shaped crystals of CG6765^1-133^, which however diffracted to above 6Å, hampering structure solution.

Hence, we used single-particle cryo-EM to elucidate the structure of BTB domain assemblies. To enhance particle contrast on cryo-EM images, we used the BTB domain of CG6765^1-133^ fused to MBP (40 kDa), which resulted in a monomer Mw of 56 kDa. Iterative masked refinement excluding flexible MBP regions resulted in a final reconstruction of core CG6765^1-133^ multimer at a resolution of 3.3 Å (full data processing flowchart is shown at Supplementary Figure S1). The map clearly shows the hexameric assembly consisting of three dimers (Figure 3a). To model atomic structure of CG6765 hexamers, we utilized an AlphaFold multimer implementation (Evans et al., 2022). The hexamer was predicted with high confidence and despite the loop regions the model fits well into the obtained cryo-EM map (Figure 3b). CG6765 hexamer consists of three canonical BTB dimers with extensive hydrophobic molecular contacts forming a β-sheet between two parallel β4-strands (corresponding to B3 according to Stogios et al (Stogios et al., 2005); Figure 3b, 3c). Interdimeric interactions were strengthened by contacts between A2 and the B1/B3 sheet, with the B3 strand formed by residues highly conserved only within the TTK group (Figure 2a). The N-terminal TTK-motif is located within the first β-strand and is involved only in dimer stabilization (Figure 3b). Central part of the hexamer forms a pore, residues of loop regions surrounding the pore are not conserved, thus it is unlikely that it can possess some physiological function.

**Figure 3.**
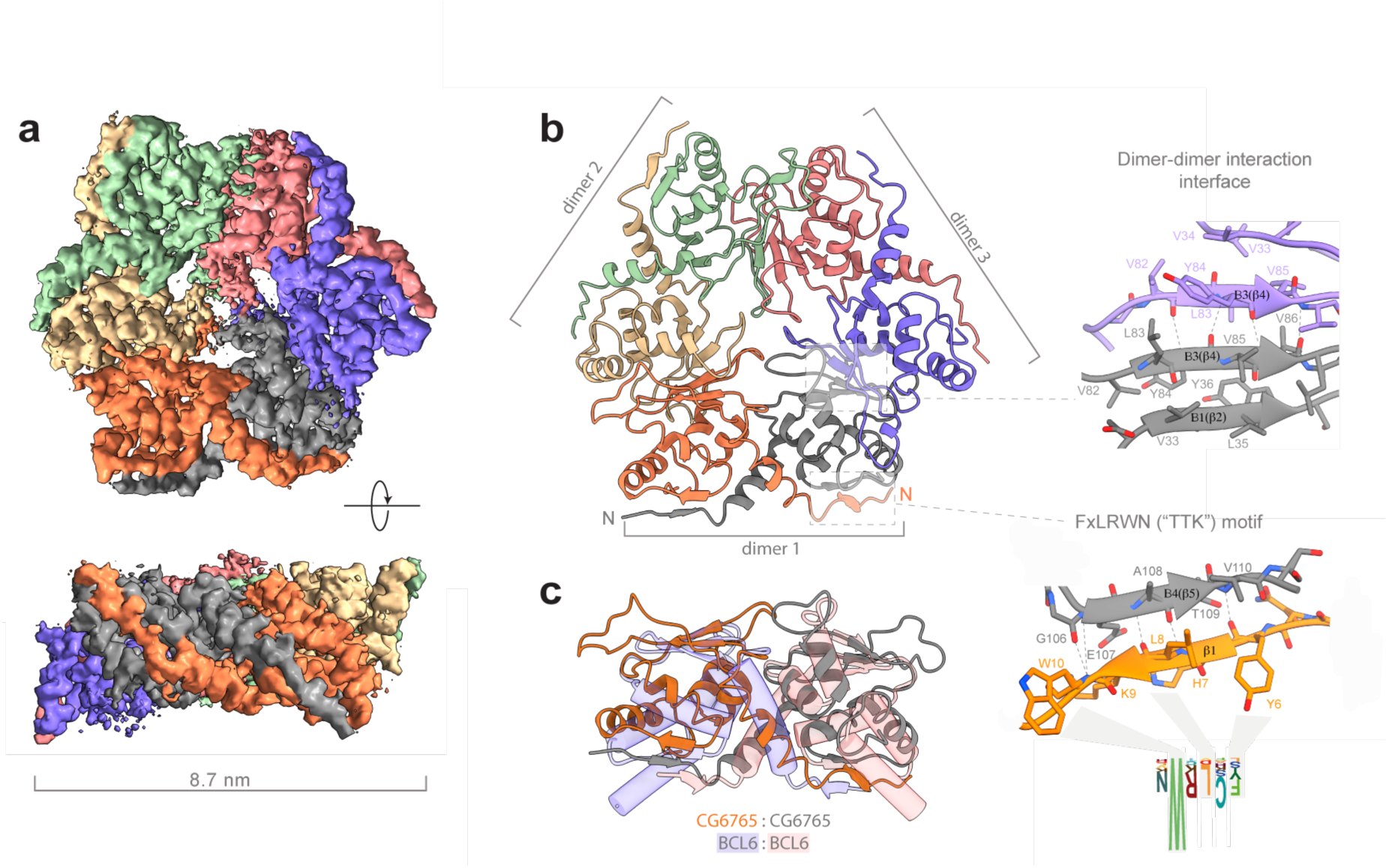
Cryo-EM structure of CG6765 BTB domain. **a** Cryo-EM map of CG6765 BTB domain. Map regions are colored according to corresponding protein chains. **b** Refined model of CG6765 BTB domain hexamer. Individual dimers are depicted, details of dimer-dimer interaction interface and beta-sheet formation by FxLRWN (“TTK”) motif are shown at the right. Secondary structure elements are depicted according to (Stogios et al., 2005). **c** Overlay of dimeric subunit of CG6765 BTB domain and classical dimer of Bcl6 BTB domain (PDB ID: 1R28) (Ahmad et al., 2003).

Thus, the structure of CG6765 BTB domain obtained with cryo-EM reveals the hexameric structure of BTB domain of TTK group consisted of three canonical BTB dimers connected via an novel interface formed by two parallel β-strands which has not yet been implicated in BTB multimerization.

### Hexamers are the main oligomeric state of TTK-type BTB domains in solution

As it was noted earlier, 8 out of 10 soluble BTB domains of TTK group formed stable high order multimers. The SEC profile for the representative BTB domain of LOLA had a single symmetric peak, the position of which remained unchanged even upon 10-fold dilution (Figure 4a). Since the apparent Mw of 113 kDa determined relative to protein standards did not allow us to unequivocally determine the oligomeric state of LOLA (the predicted monomer mass is 15.25 kDa), it was analyzed using SEC-MALS, which provides the absolute Mw. The Mw determined by this method was 86.3 kDa, which was 5.7 times larger than the predicted monomeric Mw, and therefore was closest to a hexameric species. The monodispersity of the sample (Mw/Mn = 1.000) and the unchanged position on the elution profile upon dilution together indicated that the observed oligomer is stable. Consistent with this, we observed that MBP-fused LOLA also had a single symmetrical peak on the SEC profile, and its MALS-derived Mw of 315 kDa suggested a monodisperse (Mw/Mn=1.000) ∼5.6-mer (Figure 4b).

**Figure 4.**
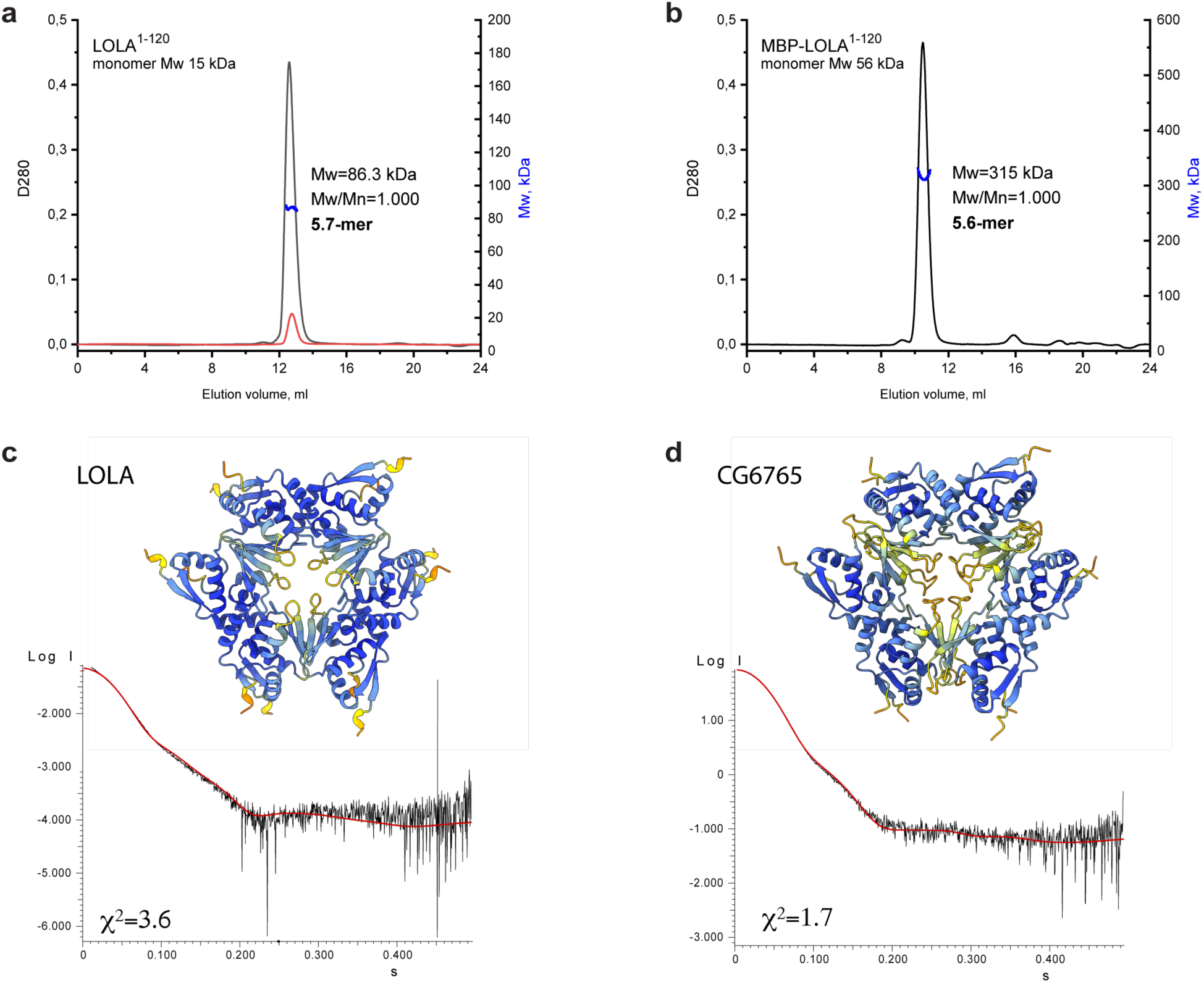
An integrative biology approach reveals the hexameric assembly of BTB domains. SEC-MALS data for LOLA **a** and MBP-LOLA **b** showing the chromatographic peaks with the Mw distributions across each peak. Average Mw values in kDa, polydispersity index (Mw/Mn) and the formally calculated oligomeric state are shown. D280 designates optical density at 280 nm (absorbance units). The second Y axes are Mw, kDa. Note that tenfold dilution of LOLA (black and red curves in panel A) did not cause any shift of the chromatographic peak, indicating stability of the observed oligomer. Alphafold2-derived models of LOLA **c** and CG6765 **d** oligomers and fits of their theoretical scattering data to the experimental SAXS data. Models are colored according to AlphaFold pLDDT values.

Since particles of BTB domains of CG6765 and LOLA are monodisperse in solution, we expected that BTB domain of LOLA will have the similar structure as was revealed by cryo-EM for CG6765. To further verify the structures and stability of stoichiometry of hexameric assemblies for LOLA BTB domain devoid of additional tags, we applied SAXS method and AlphaFold modeling using CG6765 BTB as a reference. SAXS-derived structural parameters are listed in Table 1. Estimated molecular weights for LOLA and CG6765 BTB domains roughly correspond to hexamers and are in agreement with SEC-MALS and cryo-EM data. Then, we critically assessed AlphaFold models with different stoichiometries (from dimers to octamers) using approximation of experimental SAXS data with curves calculated from the models using CRYSOL (Svergun et al., 1995) (Supplementary Figure S6a, S6b). For both CG6765 and LOLA BTBs, the theoretical scattering for hexameric models agreed best with the experimental data (χ^2^ values 1.7 and 3.6, respectively (Figure 4c, 4d)), while the fits from alternative oligomeric assemblies predicted by AlphaFold were inadequate (Supplementary Figure S6a, S6b). The models of 7-mers contained largely disconnected hexamers and an additional subunit and were not considered. The models of 5-mers provided second-best fits to the experimental SAXS profiles, but were disregarded due to symmetry considerations, contradiction to cryo-EM data and also because they appeared to represent hexamers lacking one of the subunits. We believe that, while the presence of incomplete assemblies *in vitro* cannot be excluded, they do not represent the main oligomeric state of TTK-group BTB domains. Thus, SAXS data support that the prevalent oligomer state in solution of LOLA and CG6765 BTB domains is the hexamer, in accordance with the CryoEM data. The overall structures of both hexamers are highly similar (Figure 4c, 4d).

**Table 1.** SAXS-derived structural parameters for CG6765 and LOLA BTB domains. Rg is radius of gyration, Dmax – maximum dimension of the particles, Vp – Porod volume – volume of the particles. The sample of CG6765^1-133^ with concentration of 1.5 mg/ml provided sufficiently high-quality scattering data, which were used for all fitting experiments, whereas data obtained for LOLA^1-120^ at concentrations 1.0 and 3.0 mg/ml were merged to obtain the necessary quality. Samples with higher concentrations exhibited signs of aggregation and were excluded from analysis.

| Polypeptide | Sample concentration, mg/ml | Rg, nm | Dmax, nm | Vp, nm<sup>3</sup> | Estimated molecular weight, kDa | Molecular weight of the monomer, kDa |
| --- | --- | --- | --- | --- | --- | --- |
| CG6765<sup>1-133</sup> | 1.5 | 3.7 | 12.7 | 166 | 90.5-116.5 | 15.3 |
| LOLA<sup>1-120</sup> | merged data for 1.0 and 3.0 mg/ml | 4.1 | 14.5 | 189 | 108-136 | 15.2 |

We also modelled hexamer assemblies of different BTB domains of TTK group (Supplementary Figure S7aA) including Mod(mdg4) protein (Supplementary Figure S7b), which is the best studied protein of the family so far (Golovnin et al., 2007; Melnikova et al., 2017). All models had an architecture similar to those of the LOLA and CG6765 hexamers with interdimeric interface predicted with high confidence (Supplementary Figure S7a). The dimer-dimer interaction interface is also conserved in Batman, which did not form multimers larger than dimer, according to SEC data. We therefore modelled a Batman BTB dimer and compared its structure with a LOLA dimer within the hexamer. We found no steric hindrance obstructing hexamer formation in Batman, although some hexamer-stabilizing interactions were absent in this case, for example F42, which stabilized the LOLA hexamer, is T42 in Batman, in the cryo-EM structure of the CG6765 BTB this interaction is supported by Y36 from adjacent beta-strand (Supplementary Figure S7c). From this observation, we suggest that the core β-sheet might be not sufficient for stable hexamer formation, and further stabilizing contacts are required, which may determine the specificity of the interaction. The dimer-dimer interaction interface was substantially different in the model of Chinmo BTB, which can still form multimers, as well as in Ribbon and CG15812. Unfortunately, we could not test the multimerization of the latter two domains due to strong aggregation; however, the AlphaFold model suggested that the β-strand involved in the dimer-dimer interaction remained unchanged, but confidence of prediction was substantially lower (Supplementary Figure S7a).

It is likely that unusually broad interactions between different TTK-type BTB domains (heteromeric interactions) occur due to the association of different homodimers through the interdimeric interfaces described above, which are highly similar in different members of this family (Figure 2a). We developed a set of substitutions of conserved hydrophobic residues involved in the inter-subunit β-sheet interface. In general, all mutations led to a strong decrease in protein solubility and appearance of aggregates in the SEC profiles, suggesting a general impact on protein folding (Supplementary Figure S8, S9, S10, S11 and S12). Only a small fraction of hexamers was present, along with the appearance of a fraction of dimers of comparable intensity, indicating the effect of the tested mutations on the formation of hexamers. A large peak of proteolytic fragments was also visible, suggesting protein misfolding and degradation (see Supplementary Figure S9, S10 and S11). Such a strong impact of dimer-dimer interface mutations on protein folding and self-association does not allow to confidently study the effect of these mutations on heteromeric interactions. Mutational data support the importance of the integrity of dimer-dimer interaction interface for proper folding and stability of TTK group BTB domains.

In summary, various approaches independently confirm that hexameric assembly of three dimers is the main oligomeric state of TTK group BTB domains.

### TTK-type BTB domains are specific to Arthropoda

In *D. melanogaster*, most transcription factor BTB domains are of TTK-type. We therefore investigated when these domains emerged over the course of evolution and how widespread in animals they are. We built the Hidden Markov Model (HMM) profile “TTK motif” based on sequence alignment of TTK-type BTBs from 14 Dipteran species (Figure 2b) and performed a search within the main groups of Arthropoda and several other Metazoan species (including basal groups such as Onychophora, Tardigrada, and Nematoda). TTK-type BTB domains were not found outside of Arthropoda. The most basal clades in which they emerged were Crustaceans and Arachnoidea (Figure 5a). This agrees with a recent study which traced the origin of GAF and Mod(mdg4) insulator proteins to ancestral groups of insects related to modern *Protura* and *Plecopthera* (Pauli et al., 2016). TTK-type BTB domains underwent lineage-specific expansion in modern phylogenetic groups of insects, to the most extent in Diptera and Hymenoptera. In these species, TTK-type BTB domains almost completely replace transcription factors with classical dimeric BTB domains, as in *D. melanogaster* (Figure 5a). An ortholog search of the 11 best-studied *Drosophila* TTK-type BTB proteins revealed that the oldest proteins with TTK-type BTBs are LOLA, bric-a-brac2 and Tramtrack, for which orthologs were found in the most basal clades (Figure 5b). The same search for four non-TTK *Drosophila* BTB proteins revealed that CP190 and CG6792 (a *Drosophila* PLZF homolog) are the oldest: a CP190 ortholog was found in Nematodes and CG6792 orthologs are present in almost all higher Metazoan taxa (Figure 5c). Notably, each non-TTK *Drosophila* BTB-C2H2 protein has only one isoform, whereas many factors with a TTK-type BTB have multiple isoforms, which typically differ in their DNA-binding domain, resulting in a wide diversity of these factors.

**Figure 5.**
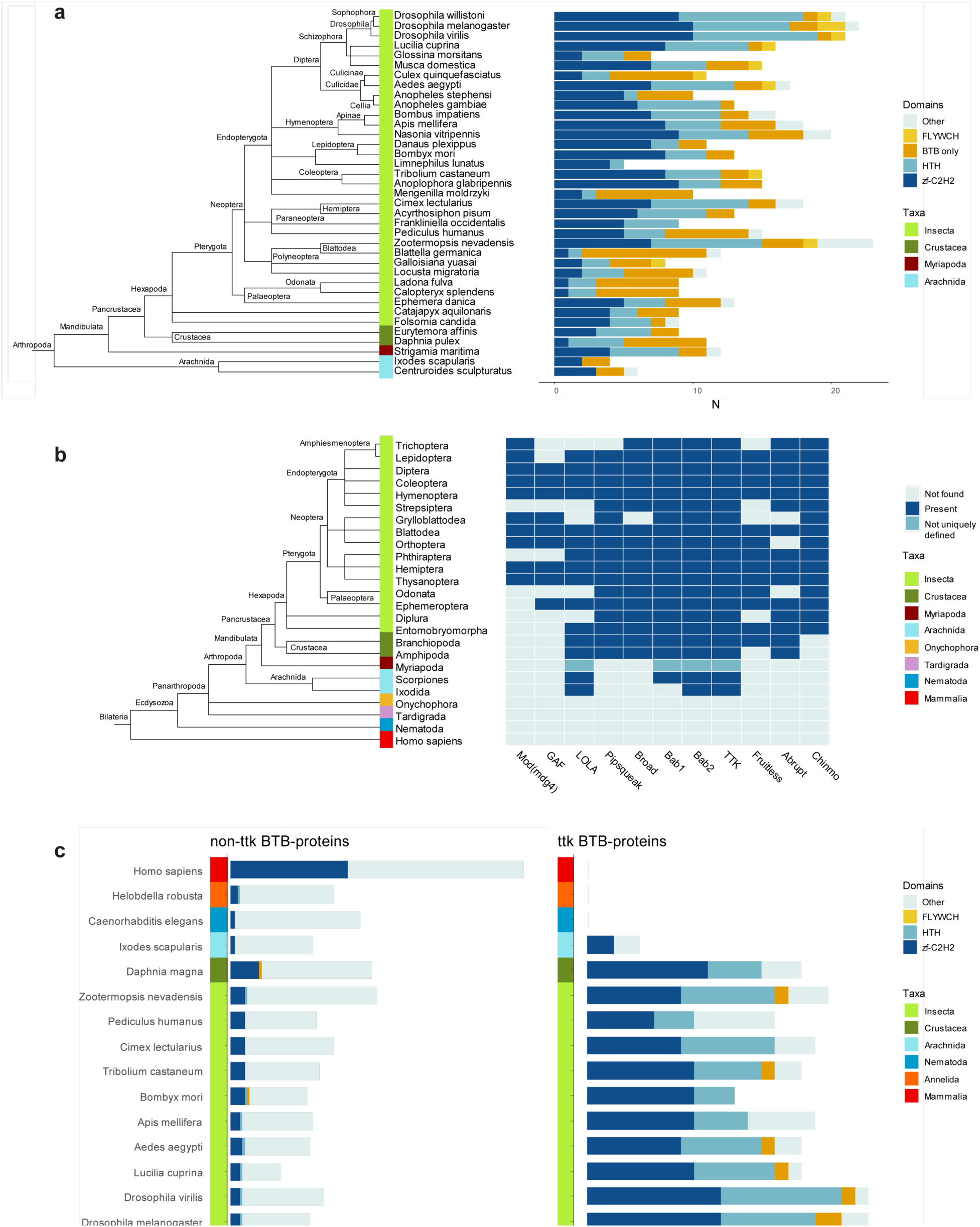
Domain architectures and orthologs of proteins with BTB domain of TTK-type in various Arthropoda lineages. **a** Phylogenetic analysis of the distribution of the main DNA and protein interaction domain types in TTK-type BTB proteins in proteomes of representatives of Hexapoda, Crustacea, Myriapoda, and Arachnida. Each type of BTB protein domain architecture is shown as a bar segment. Total search results are shown in Supplementary Table S7. The orthologs of several *Drosophila* BTB domains of TTK-type **b** and non-TTK-type **c** in proteomes of key taxa in major Arthropod and other Ecdysozoa phylogenetic groups. The phylogenetic relationships among taxa are according to NCBI Taxonomy Database. Azure blue – the ortholog is absent in the taxa, dark blue – the ortholog is present in the taxa, dark cyan – orthologs are not uniquely defined.

Most BTB proteins also contain C2H2 or HTH DNA-binding domains. With the exception of Arachnida, in all examined Arthropoda species, the FLYWCH and HTH domains are found only in combination with the TTK-type BTB domains (Figure 6a). In the human proteome, BTB-containing transcription factors almost exclusively utilize C2H2 zinc-fingers as DNA-binding domains. Interestingly, TTK-type BTBs are usually associated with one or two C2H2 domains (only MAMO has five C2H2 domains), while mammalian transcription factors commonly have BTB domains in combination with arrays consisting of an average of five C2H2 domains (Figure 6b).

**Figure 6.**
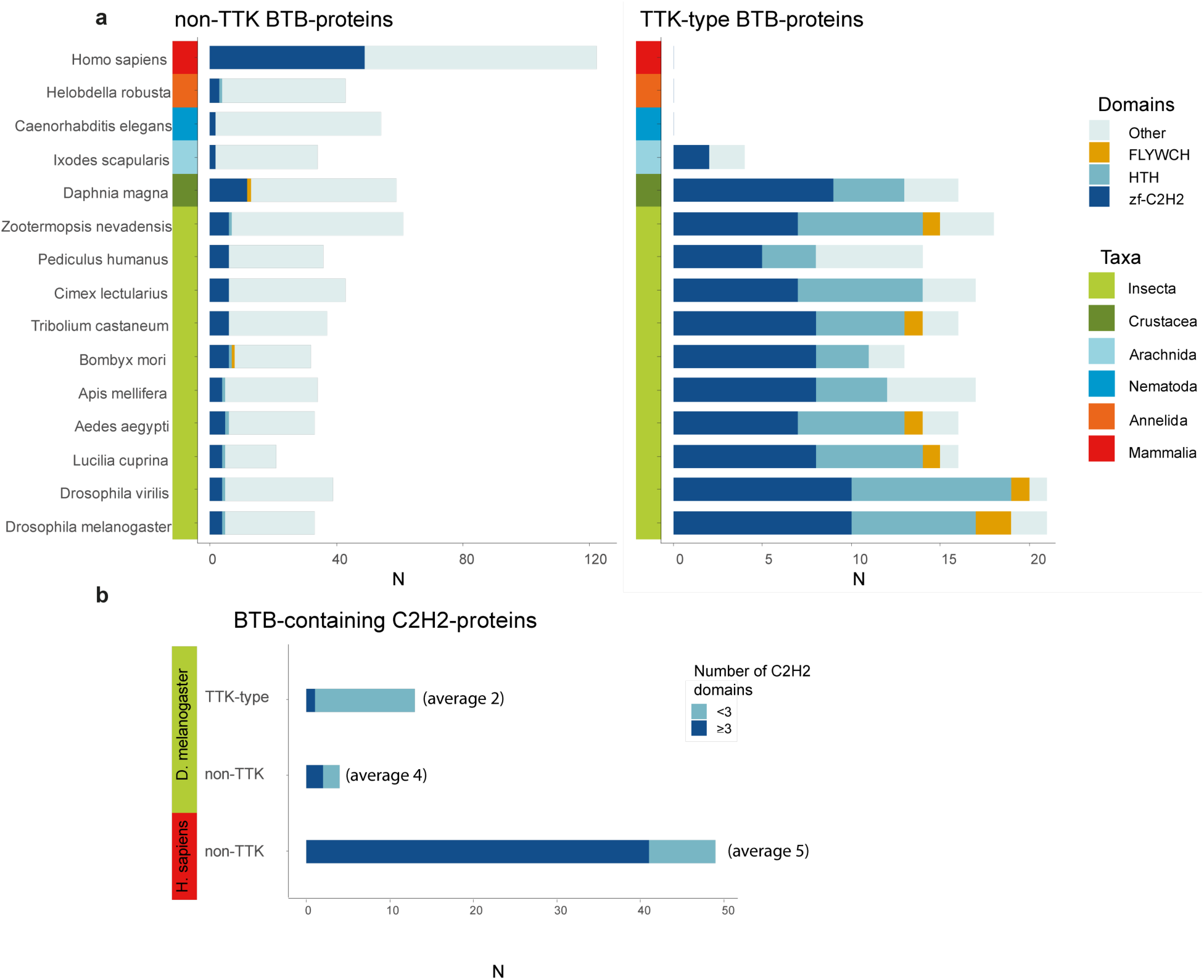
TTK-type BTB domains are specific to Arthropodan transcription factors. **a** Distribution of major DNA interaction domain types in non-TTK (left) and TTK-containing (right) BTB proteins in the proteomes of 16 Metazoan species. Each type of BTB-associated protein domain is denoted as a bar segment. Total search results are shown in Supplementary Tables S7 (TTK BTB domains) and S8 (non-TTK BTB domains). **b** Representation of non-TTK and TTK-containing BTB proteins with less than three (dark cyan) and three or more (dark blue) C2H2 domains in *Drosophila melanogaster* and *Homo sapiens* proteomes.

Taken together, our results suggest that proteins with TTK-type BTBs comprise distinct group of transcription factors specific to Arthropoda.

## Discussion

The TTK-type BTB domains form a distinct group of BTB domains specific to Arthropodan transcription factors. Here, we studied the mechanism of multimer formation of TTK-type BTB domains using an integrative structural biology approach. Cryo-EM, SAXS, molecular modeling, and MALS revealed that these domains form hexamers consisting of three dimers assembled through a novel dimer-dimer interaction interface. A neighbor surface (based on the B1-strand) was recently found to be involved in the Miz-1 BTB interaction with the HUWE-1 protein (Orth et al., 2021) and along with B3 was earlier implicated in Miz-1 tetramer formation in crystal (Stead et al., 2007). Apparently, this interface can be widely used for BTB-domain-mediated protein-protein interactions. Notably, the dimer-dimer interaction in Miz-1 involves conformational flexibility in this region, and monomers within each dimer are non-equivalent, with only one containing the B3-strand. Our structure and models show that dimers of TTK-type BTB domains consist of identical monomers; however, further structural studies will be required to elucidate precise details of this interface. Single amino acid substitutions further confirmed that hydrophobic residues at the dimer-dimer interface are critical for multimer formation. The characteristic conserved N-terminal sequence FxLRWN forms the first β-strand and was found to be important for specific dimer formation rather than for higher-order oligomerization as previously suggested (Bonchuk et al., 2011). This indicates that the FxLRWN motif at the N-terminus co-evolved together with the multimerization motif of the interface forming residues between the dimers. Probably FxLRWN is involved in binding to the yet unknown interaction partner. TTK-type BTB domains possess an unusually wide potential for heteromeric interactions despite a rather low sequence similarity. Heteromultimerization likely occurs through the interaction of different dimers, since the dimer-dimer interface is highly similar in different TTK-type BTB domains. This is a unique property of this class of BTB domains, as classical BTB domains form almost exclusively homodimers.

The expansion of the TTK-type BTB domains in insects suggests an important but poorly-understood functional role for their ability to form homo- and hetero-multimers. Most of the transcription factors of the TTK group bind to short degenerate DNA motifs. Thus, the multimerization of proteins via the BTB domain can increase the binding affinity for several motifs located in close vicinity. Such a mechanism was previously demonstrated for the pioneer and bookmarking protein GAF that is stably associated with chromatin (Bellec et al., 2022; Gaskill et al., 2021). GAF has only one zinc finger domain that binds to a GAGAG motif. It was shown that BTB oligomerization mediates strong co-operative binding of GAF to multiple sites but inhibits binding to a single motif (Espinas et al., 1999; Katsani et al., 1999).

Heteromultimerization of BTB domains can allow the formation of complexes of several TTK-type proteins on chromatin. The most striking example is the Batman protein, which consists only of the BTB domain (Mishra et al., 2003). Batman forms complexes with Pipsqueak and GAF by interacting with their BTBs (Faucheux et al., 2003). Batman also interacts with the BAB1 BTB domain, and these proteins cooperate in the control of sex combs on male tarsa (Gibert et al., 2007). Since the Batman BTB can be recruited to chromatin only through interaction with other DNA-binding BTB proteins, it does not form homomultimers that would prevent it from interaction with other BTBs. Thus, TTK-type BTB transcription factors may constitute a regulatory network controlling common target genes.

In this study, we have shown that TTK-type BTB domains are Arthropod-specific and underwent lineage-specific expansion in modern insects. The *Drosophila* proteome contains 24 transcription factors with TTK-type BTB domains. Moreover, non-TTK-type BTB domains are found in only four *Drosophila* transcription factors, suggesting that the TTK-type BTB domain confers some evolutionary-important benefits. The most basal clades in which TTK-type BTBs can be traced are Crustacea and Arachnoidea. The only such proteins present in both taxa are Tramtrack orthologs, suggesting that it is the oldest member of the family.

In human and mouse, BTB-C2H2 proteins are encoded by at least 49 genes that are important regulators of development and commonly function as sequence-specific repressors of gene expression. In contrast to Arthropoda TTK-type BTB proteins, mammalian BTB-C2H2 proteins usually have many C2H2 domains that recognize extended and specific DNA motifs. Despite the fact that mammals have more highly-specific DNA-binding proteins, many of the TTK-type BTB transcription factors in Arthropods have multiple isoforms with different DNA-binding or protein interaction domains which, along with the ability to heteromultimerize, can result in a large diversity of complexes possessing a broad range of DNA-binding specificities.

In conclusion, TTK-type BTB proteins form a structurally and functionally distinct group of Arthropod key regulatory factors with unique functions in the process of cell differentiation and transcriptional regulation. What functional role is played by the ability of these transcription factors to form various combinations of heterologous complexes remains an open and very interesting question.

## Materials and methods

### Bioinformatics

The search for orthologs for 24 *D. melanogaster* proteins with BTB domains containing ttk sequences was carried out in Metazoa, with the exception of Chordata, using the OrthoDB database (Kriventseva et al., 2019). There were no records for Mod(mdg4), GAF, and LOLA in OrthoDB, and orthologs of Fruitless were found only in Diptera. In order to fill up the data obtained from OrthoDB, orthologs for these proteins were searched in Ensembl using biomaRt (Durinck et al., 2009). The total number of identified orthologs was 3027. Amino acid sequences of orthologs of 23 ttk proteins of *D. melanogaster* in 14 Diptera species (five *Drosophila* species, two *Aedes* species, two *Anopheles* species, one *Culex* and fore flies) were aligned in MUSCLE (Edgar, 2004). The motif of the ttk domain in the resulting alignment was isolated using the EM algorithm by positional weight matrix (PWM) (at no limit motif E-value threshold and motif size 6-10 aa (Figure 2a)) in MEME (MEME SUITE) (Bailey et al., 2009), then in FIMO (MEME SUITE) we searched for the motif in the obtained database of orthologous sequences for all Metazoa (the threshold FDR value was 0.001). In addition, for the training set for 14 Diptera species, an hmm profile for the ttk domain was generated in MEGA-X (Kumar et al., 2018), and it was searched using HMMSEARCH (HmmerWeb version 2.41.1 (Potter et al., 2018)) in the UniProtKB database at E-value = 10. Data on full protein sequences (n = 259, excluding isoforms) deposited in UniProtKB were combined with previously obtained ttk-containing sequences of orthologs from OrthoDB and Ensembl (n = 2208). In addition, in the large taxa closest to Arthropoda (several phyla of Protostomes: Tardigrada, Onychophora, Priapulida, Kinorhyncha, Loricifera, Nematoda, Annelida, Mollusca), the search for the ttk sequence motif in blastp and hmm-profile in HMMSEARCH were carried out; in the representatives of the above taxa, no proteins containing the ttk motif were found in non-redundant DBs.

Phylogenetic classification was reconstructed in accordance with the NCBI Taxonomy Database using taxize (Chamberlain and Szocs, 2013), visualization of the phylogenetic tree is implemented in ggtree (Yu, 2020). In the most basal clades (Crustacea, Arachnida, Myriapoda), the presence of the corresponding orthologs was predicted manually in blastp, due to the low similarity of amino acid sequences. We considered as orthologs, in addition to the previously annotated ones, only those proteins for which homology was observed in the structure, in both the BTB domain (including the presence of the ttk sequence) and the DNA binding domain. The presence/absence of some orthologs in individual orders of insects was also checked in blastp and HMMSEARCH (HmmerWeb version 2.41.1). In addition, the previously published data by Pauli et al. (Pauli et al., 2016) were used for Mod (mdg4) and GAF.

The domain structure of the detected orthologs of ttk-containing proteins was studied in 37 reference species - representatives of 16 Hexapoda orders, two orders of Crustaceans and Arachnids, and one - Myriapoda (Figure 5a). The search for domains in the orthologous sequences was performed using the Pfam database in Batch CD-Search (CDD / SPARCLE NCBI (Lu, Wang et al., 2020)), the threshold E-value was taken to be 0.5. The detection results were filtered: incomplete and overlapping domains were removed. In sum, after filtering, the total number of domains found was 1055; the analyzed domains belonged to 26 PFAM families.

For 16 species with well-annotated proteomes belonging to the phyla Arthropoda, Nematoda, Annelida, as well as for *Homo sapiens*, the sequences of all proteins containing BTB domains in their proteomes were searched using InterProScan in the Pfam protein domain database by the generalized hmm -BTB domain profile. In the set of the found sequences, the search for the ttk-domain motif in FIMO was carried out as described above, the ttk-containing proteins were filtered, and the remaining sequences were analyzed in Batch CD-Search in order to identify their domain structure. In total, 3028 domains belonging to 99 PFAM families were found in the sequences of BTB proteins without the ttk motif in 16 species.

### Plasmids and cloning

cDNAs of BTB domains were PCR-amplified using corresponding primers (Supplementary Table S1) and cloned into modified pET32a(+) vector (Novagen) encoding TEV protease cleavage site after 6xHis-tag and Thioredoxin, and into pGAD424 and pGBT9 vectors (Clontech) in frame with GAL4 Activation or DNA-binding domains, respectively. PCR-directed mutagenesis was used to create constructs expressing mutant BTBs using mutagenic primers (Supplementary Table S1). For MBP fusions cDNAs of BTB domains were cloned into pMALX(A) vector (Moon et al., 2010).

### Protein expression and purification

BL21(DE3) cells transformed with a construct expressing BTB domain fused with TEV-cleavable 6xHis-Thioredoxin were grown in 1 L of LB media to an D600 of 1.0 at 37 °C and then induced with 1 mM IPTG at 18 °C overnight. Cells were disrupted by sonication in buffer A (30 mM HEPES (pH 7.5), 400 mM NaCl, 5 mM β-mercaptoethanol, 5% glycerol, 0.1% NP40, 10 mM imidazole) containing 1 mM PMSF and Calbiochem Complete Protease Inhibitor Cocktail VII (1 μL/ml). After centrifugation, lysate was applied to a Ni-NTA column, and after washing with buffer B (30 mM HEPES (pH 7.5), 400 mM NaCl, 5 mM β-mercaptoethanol, 30 mM imidazole) was eluted with 300 mM imidazole. For cleavage of the 6x-His-thioredoxin-tag, 6x-His-tagged TEV protease was added at a molar ratio of 1:50 directly to the eluted protein and the mixture was incubated for 2 hours at room temperature, then dialyzed against buffer A without NP-40 and applied to a Ni-NTA column. Flow-through was collected; dialyzed against 20 mM Tris-HCl (pH 7.4), 50 mM NaCl and 1 mM DTT; and then applied to a SOURCE15Q 4.6/100 column (GE Healthcare) and eluted with a 50-500 mM linear gradient of NaCl. Fractions containing protein were concentrated, frozen in liquid nitrogen, and stored at -70 °C. Size-exclusion chromatography was performed in 20 mM Tris-HCl (pH 7.4), 200 mM NaCl and 1 mM DTT using Superdex S200 10/300GL column (GE Healthcare).

### SEC-MALS

Size-exclusion chromatography with multi-angle light scattering (SEC-MALS) detection was used to determine absolute Mw for LOLA and MBP-LOLA samples. Protein samples (1-5 mg/ml) were loaded individually on a Superdex 200 Increase 10/300 column (GE Healthcare), and the elution profiles were obtained using a tandem of sequentially connected UV-Vis Prostar 335 (Varian, Australia) and miniDAWN detectors (Wyatt Technology, USA). The column was pre-equilibrated with filtered (0.1 μm) and degassed 20 mM Tris-HCl buffer, pH 7.6, containing 200 mM NaCl and 5 mM β-mercaptoethanol and was operated at a 0.8 ml/min flow rate. Data were processed in ASTRA 8.0 (Wyatt Technology, USA) using dn/dc equal to 0.185 and extinction coefficients ε(0.1%) at 280 nm equal to 0.80 ml/(mg cm) and 1.38 ml/(mg cm) for LOLA and MBP-LOLA, respectively. Additionally, apparent Mw values were determined from column calibration with standard proteins. Data were processed and presented using Origin 9.0 (Originlab, Northampton, MA, USA).

### Cryo-EM grid preparation and data collection

Quantifoil R1.2/1.3 300-mesh carbon grids were glow-discharged for 60 s at 20 mA using a GloQube (Quorum) instrument. 4 μl of the MBP-CG6765^1-133^ sample (4 μg/μl) were applied to the freshly glow-discharged grids and plunge-frozen in LN2-cooled liquid ethane using a Vitrobot Mark IV (Thermo Fisher Scientific) with a blotting time of 2.5 s. Temperature and relative humidity were maintained at 4 °C and 100%, respectively. Grids were clipped and loaded into a 300-kV Titan Krios G4i microscope (Thermo Fisher Scientific, USA) equipped with a Selectris X energy filter and a Falcon 4i (Thermo Fisher Scientific, USA) direct electron detector. Micrographs were recorded at a nominal magnification of ×165,000 corresponding to a calibrated pixel size of 0.729 Å. A total of 9765 movies were recorded with a total dose of ∼40 electron/Å2 per movie. Defocus range was set between −0.5 μm and −2 μm. Data collection and processing statistics are summarized in Supplementary Table S2.

### Cryo-EM data processing

Supplementary Figure S1 illustrates the data processing workflow for the MBP-CG6765^1-133^ dataset. The following pre-processing steps were performed with cryoSPARC Live v4.3.1 (Punjani et al., 2017). Movie stacks were motion-corrected and dose-weighted, contrast transfer function (CTF) estimates for the motion-corrected micrographs were calculated. Particles were initially picked with a blob-picker using subset-selected micrographs, and these were used for reference-free two-dimensional (2D) classification to generate picking templates. Subsequent image processing was carried out with cryoSPARC v4.3.1 (Punjani et al., 2017). The templates were used to train a picking model in Topaz (Bepler et al., 2019), which was subsequently used to pick particles from the whole dataset. Auto-picking using Topaz from 9765 micrographs yielded ∼100000 particles. Initial models were generated without imposing symmetry (C1) using stochastic gradient descent in cryoSPARC Live. Best initial model was subjected to homogeneous and heterogeneous refinement rounds and best classes were used to create templates for another round of template picking, followed by 2D classification and training new Topaz picking model, which was used for final picking round. 480721 particles were picked. Particles were classified with three-dimensional (3D) heterogenous refinement using four classes, resulting in 197,562 particles.

To generate a high-resolution reconstruction, particles were re-extracted in 448 pixels box size followed by 3D refinement with local angular search using the map from the previous processing in cryoSPARC as reference. A mask including only map fragments corresponding to hexamer core was generated using UCSF Chimera (Pettersen et al., 2004) and RELION v.4.0 (Scheres, 2012). The mask was used in subsequent 3D refinement jobs and CTF refinements in CryoSPARC. The particles were exported to RELION using PyEM (Asarnow et al., 2019) and further processed in RELION 4.01. After 3D refinement followed by CTF refinement, Bayesian polishing was applied. 3D refinement on the polished particles, followed by CTF refinement and another round of 3D refinement with Blush regularization (in RELION 5.0beta (Kimanius et al., 2023)), yielded a reconstruction to ∼3.3 Å overall resolution with C1 symmetry. We realized that Blush regularization did not improve the nominal resolution, however it slightly improved the quality of the map especially in the core region of the complex. In the peripheral solvent exposed part, the map without blush regularization appeared to be of better quality and was therefore used to build the loop regions of the complex. Both maps were deposited in the EMDB.

AlphaFold model of the hexamer was used as initial model, it was fitted into the reconstruction using UCSF Chimera (Pettersen et al., 2004), followed by manual real-space refinement in Coot, and further refined with Phenix.refine (Liebschner et al., 2019) and ISOLDE (Croll, 2018).

### Yeast two-hybrid assay

Yeast two-hybrid assay was performed as described (Bonchuk et al., 2011).

### SAXS data collection and analysis

Synchrotron radiation X-ray scattering data were collected using standard procedures on the BM29 BioSAXS beamline at the ESRF (Grenoble, France) as described previously (Bonchuk et al., 2020). Data analysis was performed using ATSAS software package (Franke et al., 2017). Approximation of the experimental scattering profiles using calculated scattering curves was performed with CRYSOL (Svergun et al., 1995). The molecular mass (MM) of the protein was calculated using several algorithms implemented in ATSAS package (Franke et al., 2017).

## Supporting information

Models of BTB domain hexamers and SAXS fits

## Acknowledgements

This work was supported by the Russian Science Foundation – project 19-74-10099-P to A.B. (expression and purification of proteins and their mutants), project 19-74-30026-Р to P.G. (analysis of protein-protein interactions) and by Ministry of Science and Higher Education of the Russian Federation — grant 075-15-2019-1661 (structural and bioinformatic analysis). Funding for open access charge: Ministry of Science and Higher Education of the Russian Federation and Russian Science Foundation. The single-particle cryo-EM work was financially supported by the KAUST Baseline Grant BAS/1/1107-01-01. N.N.S. and K.M.B. acknowledges that SEC-MALS work was supported by the Ministry of Science and Higher Education of the Russian Federation. We are grateful to Dr. Alexander Kuklin (Joint Institute of Nuclear Research, Dubna) for help in SAXS data collection. We acknowledge the European Synchrotron Radiation Facility for provision of synchrotron radiation facilities and we would like to thank the staff of the ESRF for assistance and support in using beamline BM29. We thank Andrey Moiseenko (MSU, Moscow, Russia) for his help in negative-staining EM data collection.

## Author contributions

A.N.B. initiated the project, designed the research studies, gathered the research team, supervised the experiments, conducted protein expression and purification experiments, acquired SAXS data, analyzed cryo-EM data, prepared figures and drafted, wrote, and revised the manuscript. K.I.B. conducted protein interaction assays. R.B. conducted cryo-EM experiments and acquired data. K.M.B. conducted crystallization trials and molecular modelling. N.N.S. conducted SEC-MALS experiments, and analyzed SAXS data. A.D.B. conducted negative staining electron microscopy experiments. O.V.A. conducted protein expression and purification experiments. A.M.K. conducted bioinformatic analysis. K.Y.K. conducted protein interaction assays. V.O.P. designed the research studies, supervised the experiments. A.N. analyzed cryo-EM data, refined the molecular model, and revised the manuscript. P.G.G. designed the research studies, supervised the experiments, and revised the manuscript.

## Data availability

The cryo-EM maps (blushed-regularized and normal regularization) and PDB model files have been deposited in the Protein Data Bank under the PDB entry code 8RC6 and in the EMDB with entry code EMD-19049. SAXS data have been deposited in the Small Angle Scattering Biological Data Bank (www.sasbdb.org) under accession codes SASDP59 (merged data for LOLA^1-120^ at 1.0 mg/ml and 3.0 mg/ml), SASDP49 (CG6765^1-133^ at 1.5 mg/ml). Atomic models (both native and exactly corresponding to expression constructs) and reports of SAXS approximation are provided as Supplemental files. Results of bioinformatic analysis are provided as Supplemental tables.

## Supplementary Data

Supplementary Data are available online.

## References

1. Ahmad, K.F., Engel, C.K., and Prive, G.G. 1998. Crystal structure of the BTB domain from PLZF. Proc Natl Acad Sci U S A 95: 12123–12128. DOl: 10.1073/pnas.95.21.12123

2. Ahmad, K.F., Melnick, A., Lax, S., Bouchard, D., Liu, J., Kiang, C.L., Mayer, S., Takahashi, S., Licht, J.D., and Prive, G.G. 2003. Mechanism of SMRT corepressor recruitment by the BCL6 BTB domain. Mol Cell 12: 1551–1564. DOl: 10.1016/s1097-2765(03)00454-4

3. Asarnow, D., Palovcak, E., and Cheng, Y. 2019. UCSF pyem v0.5. Zenodo 10.5281/zenodo.3576630. DOl:

4. Bailey, T.L., Boden, M., Buske, F.A., Frith, M., Grant, C.E., Clementi, L., Ren, J., Li, W.W., and Noble, W.S. 2009. MEME SUITE: tools for motif discovery and searching. Nucleic Acids Res 37: W202–208. DOl: 10.1093/nar/gkp335

5. Bartkuhn, M., Straub, T., Herold, M., Herrmann, M., Rathke, C., Saumweber, H., Gilfillan, G.D., Becker, P.B., and Renkawitz, R. 2009. Active promoters and insulators are marked by the centrosomal protein 190. EMBO J 28: 877–888. DOl: 10.1038/emboj.2009.34

6. Beaster-Jones, L., and Okkema, P.G. 2004. DNA binding and in vivo function of C.elegans PEB-1 require a conserved FLYWCH motif. J Mol Biol 339: 695–706. DOl: 10.1016/j.jmb.2004.04.030

7. Bellec, M., Dufourt, J., Hunt, G., Lenden-Hasse, H., Trullo, A., Zine El Aabidine, A., Lamarque, M., Gaskill, M.M., Faure-Gautron, H., Mannervik, M., Harrison, M.M., Andrau, J.C., Favard, C., Radulescu, O., and Lagha, M. 2022. The control of transcriptional memory by stable mitotic bookmarking. Nat Commun 13: 1176. DOl: 10.1038/s41467-022-28855-y

8. Bepler, T., Morin, A., Rapp, M., Brasch, J., Shapiro, L., Noble, A.J., and Berger, B. 2019. Positive-unlabeled convolutional neural networks for particle picking in cryo-electron micrographs. Nat Methods 16: 1153–1160. DOl: 10.1038/s41592-019-0575-8

9. Bertolini, M., Fenzl, K., Kats, I., Wruck, F., Tippmann, F., Schmitt, J., Auburger, J.J., Tans, S., Bukau, B., and Kramer, G. 2021. Interactions between nascent proteins translated by adjacent ribosomes drive homomer assembly. Science 371: 57–64. DOl: 10.1126/science.abc7151

10. Bonchuk, A., Balagurov, K., and Georgiev, P. 2023. BTB domains: A structural view of evolution, multimerization, and protein-protein interactions. Bioessays 45: e2200179. DOl: 10.1002/bies.202200179

11. Bonchuk, A., Denisov, S., Georgiev, P., and Maksimenko, O. 2011. Drosophila BTB/POZ domains of “ttk group” can form multimers and selectively interact with each other. J Mol Biol 412: 423–436. DOl: 10.1016/j.jmb.2011.07.052

12. Bonchuk, A., Kamalyan, S., Mariasina, S., Boyko, K., Popov, V., Maksimenko, O., and Georgiev, P. 2020. N-terminal domain of the architectural protein CTCF has similar structural organization and ability to self-association in bilaterian organisms. Sci Rep 10: 2677. DOl: 10.1038/s41598-020-59459-5

13. Bradley, P.L., and Andrew, D.J. 2001. ribbon encodes a novel BTB/POZ protein required for directed cell migration in Drosophila melanogaster. Development 128: 3001–3015. DOl:

14. Buchner, K., Roth, P., Schotta, G., Krauss, V., Saumweber, H., Reuter, G., and Dorn, R. 2000. Genetic and molecular complexity of the position effect variegation modifier mod(mdg4) in Drosophila. Genetics 155: 141–157. DOl:

15. Chaharbakhshi, E., and Jemc, J.C. 2016. Broad-complex, tramtrack, and bric-a-brac (BTB) proteins: Critical regulators of development. Genesis (New York, NY : 2000) 54: 505–518. DOl: 10.1002/dvg.22964

16. Chamberlain, S.A., and Szocs, E. 2013. taxize: taxonomic search and retrieval in R. F1000Res 2: 191. DOl: 10.12688/f1000research.2-191.v2

17. Croll, T.I. 2018. ISOLDE: a physically realistic environment for model building into low-resolution electron-density maps. Acta Crystallogr D Struct Biol 74: 519–530. DOl: 10.1107/S2059798318002425

18. Durinck, S., Spellman, P.T., Birney, E., and Huber, W. 2009. Mapping identifiers for the integration of genomic datasets with the R/Bioconductor package biomaRt. Nat Protoc 4: 1184–1191. DOl: 10.1038/nprot.2009.97

19. Edgar, R.C. 2004. MUSCLE: multiple sequence alignment with high accuracy and high throughput. Nucleic Acids Res 32: 1792–1797. DOl: 10.1093/nar/gkh340

20. Espinas, M.L., Jimenez-Garcia, E., Vaquero, A., Canudas, S., Bernues, J., and Azorin, F. 1999. The N-terminal POZ domain of GAGA mediates the formation of oligomers that bind DNA with high affinity and specificity. J Biol Chem 274: 16461–16469. DOl:

21. Evans, R., O’Neill, M., Pritzel, A., Antropova, N., Senior, A., Green, T., Žídek, A., Bates, R., Blackwell, S., Yim, J., Ronneberger, O., Bodenstein, S., Zielinski, M., Bridgland, A., Potapenko, A., Cowie, A., Tunyasuvunakool, K., Jain, R., Clancy, E., Kohli, P., Jumper, J., and Hassabis, D. 2022. Protein complex prediction with AlphaFold-Multimer. bioRxiv https://www.biorxivorg/content/101101/20211004463034v2. DOl: 10.1101/2021.10.04.463034

22. Faucheux, M., Roignant, J.Y., Netter, S., Charollais, J., Antoniewski, C., and Theodore, L. 2003. batman Interacts with polycomb and trithorax group genes and encodes a BTB/POZ protein that is included in a complex containing GAGA factor. Mol Cell Biol 23: 1181–1195. DOl: 10.1128/MCB.23.4.1181-1195.2003

23. Franke, D., Petoukhov, M.V., Konarev, P.V., Panjkovich, A., Tuukkanen, A., Mertens, H.D.T., Kikhney, A.G., Hajizadeh, N.R., Franklin, J.M., Jeffries, C.M., and Svergun, D.I. 2017. ATSAS 2.8: a comprehensive data analysis suite for small-angle scattering from macromolecular solutions. J Appl Crystallogr 50: 1212–1225. DOl: 10.1107/S1600576717007786

24. Gaskill, M.M., Gibson, T.J., Larson, E.D., and Harrison, M.M. 2021. GAF is essential for zygotic genome activation and chromatin accessibility in the early Drosophila embryo. Elife 10. DOl: 10.7554/eLife.66668

25. Ghetu, A.F., Corcoran, C.M., Cerchietti, L., Bardwell, V.J., Melnick, A., and Prive, G.G. 2008. Structure of a BCOR corepressor peptide in complex with the BCL6 BTB domain dimer. Mol Cell 29: 384–391. DOl: 10.1016/j.molcel.2007.12.026S1097-2765(08)00069-5 [pii]

26. Gibert, J.M., Peronnet, F., and Schlotterer, C. 2007. Phenotypic plasticity in Drosophila pigmentation caused by temperature sensitivity of a chromatin regulator network. PLoS Genet 3: e30. DOl: 10.1371/journal.pgen.0030030

27. Golovnin, A., Mazur, A., Kopantseva, M., Kurshakova, M., Gulak, P.V., Gilmore, B., Whitfield, W.G., Geyer, P., Pirrotta, V., and Georgiev, P. 2007. Integrity of the Mod(mdg4)-67.2 BTB domain is critical to insulator function in Drosophila melanogaster. Mol Cell Biol 27: 963–974. DOl: 10.1128/MCB.00795-06

28. Horiuchi, T., Giniger, E., and Aigaki, T. 2003. Alternative trans-splicing of constant and variable exons of a Drosophila axon guidance gene, lola. Genes Dev 17: 2496–2501. DOl: 10.1101/gad.1137303

29. Huang, D.H., Chang, Y.L., Yang, C.C., Pan, I.C., and King, B. 2002. pipsqueak encodes a factor essential for sequence-specific targeting of a polycomb group protein complex. Mol Cell Biol 22: 6261–6271. DOl: 10.1128/MCB.22.17.6261-6271.2002

30. Katsani, K.R., Hajibagheri, M.A., and Verrijzer, C.P. 1999. Co-operative DNA binding by GAGA transcription factor requires the conserved BTB/POZ domain and reorganizes promoter topology. EMBO J 18: 698–708. DOl: 10.1093/emboj/18.3.698

31. Kimanius, D., Jamali, K., Wilkinson, M.E., Lövestam, S., Velazhahan, V., Nakane, T., and Scheres, S.H.W. 2023. Data-driven regularisation lowers the size barrier of cryo-EM structure determination. *bioRxiv* https://www.biorxivorg/content/101101/20231023563586v1. DOl:

32. Kriventseva, E.V., Kuznetsov, D., Tegenfeldt, F., Manni, M., Dias, R., Simao, F.A., and Zdobnov, E.M. 2019. OrthoDB v10: sampling the diversity of animal, plant, fungal, protist, bacterial and viral genomes for evolutionary and functional annotations of orthologs. Nucleic Acids Res 47: D807–D811. DOl: 10.1093/nar/gky1053

33. Kumar, S., Stecher, G., Li, M., Knyaz, C., and Tamura, K. 2018. MEGA X: Molecular Evolutionary Genetics Analysis across Computing Platforms. Mol Biol Evol 35: 1547–1549. DOl: 10.1093/molbev/msy096

34. Li, X., Peng, H., Schultz, D.C., Lopez-Guisa, J.M., Rauscher, F.J., 3rd, and Marmorstein, R. 1999. Structure-function studies of the BTB/POZ transcriptional repression domain from the promyelocytic leukemia zinc finger oncoprotein. Cancer Res 59: 5275–5282. DOl:

35. Liebschner, D., Afonine, P.V., Baker, M.L., Bunkoczi, G., Chen, V.B., Croll, T.I., Hintze, B., Hung, L.W., Jain, S., McCoy, A.J., Moriarty, N.W., Oeffner, R.D., Poon, B.K., Prisant, M.G., Read, R.J., Richardson, J.S., Richardson, D.C., Sammito, M.D., Sobolev, O.V., Stockwell, D.H., Terwilliger, T.C., Urzhumtsev, A.G., Videau, L.L., Williams, C.J., and Adams, P.D. 2019. Macromolecular structure determination using X-rays, neutrons and electrons: recent developments in Phenix. Acta Crystallogr D Struct Biol 75: 861–877. DOl: 10.1107/S2059798319011471

36. Lomaev, D., Mikhailova, A., Erokhin, M., Shaposhnikov, A.V., Moresco, J.J., Blokhina, T., Wolle, D., Aoki, T., Ryabykh, V., Yates, J.R., 3rd, Shidlovskii, Y.V., Georgiev, P., Schedl, P., and Chetverina, D. 2017. The GAGA factor regulatory network: Identification of GAGA factor associated proteins. PLoS One 12: e0173602. DOl: 10.1371/journal.pone.0173602 PONE-D-16-28049 [pii]

37. Maeng, O., Son, W., Chung, J., Lee, K.S., Lee, Y.H., Yoo, O.J., Cha, G.H., and Paik, S.G. 2012. The BTB/POZ-ZF transcription factor dPLZF is involved in Ras/ERK signaling during Drosophila wing development. Mol Cells 33: 457–463. DOl: 10.1007/s10059-012-2179-3

38. Melnikova, L., Kostyuchenko, M., Molodina, V., Parshikov, A., Georgiev, P., and Golovnin, A. 2017. Multiple interactions are involved in a highly specific association of the Mod(mdg4)-67.2 isoform with the Su(Hw) sites in Drosophila. Open Biol 7. DOl: 170150 [pii] 10.1098/rsob.170150rsob.170150 [pii]

39. Mena, E.L., Jevtic, P., Greber, B.J., Gee, C.L., Lew, B.G., Akopian, D., Nogales, E., Kuriyan, J., and Rape, M. 2020. Structural basis for dimerization quality control. Nature 586: 452–456. DOl: 10.1038/s41586-020-2636-7

40. Mena, E.L., Kjolby, R.A.S., Saxton, R.A., Werner, A., Lew, B.G., Boyle, J.M., Harland, R., and Rape, M. 2018. Dimerization quality control ensures neuronal development and survival. Science 362. DOl: 10.1126/science.aap8236

41. Mishra, K., Chopra, V.S., Srinivasan, A., and Mishra, R.K. 2003. Trl-GAGA directly interacts with lola like and both are part of the repressive complex of Polycomb group of genes. Mech Dev 120: 681–689. DOl: 10.1016/s0925-4773(03)00046-7

42. Moon, A.F., Mueller, G.A., Zhong, X., and Pedersen, L.C. 2010. A synergistic approach to protein crystallization: combination of a fixed-arm carrier with surface entropy reduction. Protein Sci 19: 901–913. DOl: 10.1002/pro.368

43. Mukai, M., Hayashi, Y., Kitadate, Y., Shigenobu, S., Arita, K., and Kobayashi, S. 2007. MAMO, a maternal BTB/POZ-Zn-finger protein enriched in germline progenitors is required for the production of functional eggs in Drosophila. Mech Dev 124: 570–583. DOl: 10.1016/j.mod.2007.05.001

44. Oliver, D., Sheehan, B., South, H., Akbari, O., and Pai, C.Y. 2010. The chromosomal association/dissociation of the chromatin insulator protein Cp190 of Drosophila melanogaster is mediated by the BTB/POZ domain and two acidic regions. BMC Cell Biol 11: 101. DOl: 10.1186/1471-2121-11-101

45. Olivieri, D., Paramanathan, S., Bardet, A.F., Hess, D., Smallwood, S.A., Elling, U., and Betschinger, J. 2021. The BTB-domain transcription factor ZBTB2 recruits chromatin remodelers and a histone chaperone during the exit from pluripotency. J Biol Chem 297: 100947. DOl: 10.1016/j.jbc.2021.100947

46. Orth, B., Sander, B., Moglich, A., Diederichs, K., Eilers, M., and Lorenz, S. 2021. Identification of an atypical interaction site in the BTB domain of the MYC-interacting zinc-finger protein 1. Structure 29: 1230–1240 e1235. DOl: 10.1016/j.str.2021.06.005

47. Pauli, T., Vedder, L., Dowling, D., Petersen, M., Meusemann, K., Donath, A., Peters, R.S., Podsiadlowski, L., Mayer, C., Liu, S., Zhou, X., Heger, P., Wiehe, T., Hering, L., Mayer, G., Misof, B., and Niehuis, O. 2016. Transcriptomic data from panarthropods shed new light on the evolution of insulator binding proteins in insects : Insect insulator proteins. BMC Genomics 17: 861. DOl: 10.1186/s12864-016-3205-1

48. Perez-Torrado, R., Yamada, D., and Defossez, P.A. 2006. Born to bind: the BTB protein-protein interaction domain. Bioessays 28: 1194–1202. DOl: 10.1002/bies.20500

49. Pettersen, E.F., Goddard, T.D., Huang, C.C., Couch, G.S., Greenblatt, D.M., Meng, E.C., and Ferrin, T.E. 2004. UCSF Chimera--a visualization system for exploratory research and analysis. J Comput Chem 25: 1605–1612. DOl: 10.1002/jcc.20084

50. Plevock, K.M., Galletta, B.J., Slep, K.C., and Rusan, N.M. 2015. Newly Characterized Region of CP190 Associates with Microtubules and Mediates Proper Spindle Morphology in Drosophila Stem Cells. PLoS One 10: e0144174. DOl: 10.1371/journal.pone.0144174

51. Potter, S.C., Luciani, A., Eddy, S.R., Park, Y., Lopez, R., and Finn, R.D. 2018. HMMER web server: 2018 update. Nucleic Acids Res 46: W200–W204. DOl: 10.1093/nar/gky448

52. Punjani, A., Rubinstein, J.L., Fleet, D.J., and Brubaker, M.A. 2017. cryoSPARC: algorithms for rapid unsupervised cryo-EM structure determination. Nat Methods 14: 290–296. DOl: 10.1038/nmeth.4169

53. Sabirov, M., Kyrchanova, O., Pokholkova, G.V., Bonchuk, A., Klimenko, N., Belova, E., Zhimulev, I.F., Maksimenko, O., and Georgiev, P. 2021a. Mechanism and functional role of the interaction between CP190 and the architectural protein Pita in Drosophila melanogaster. Epigenetics Chromatin 14: 16. DOl: 10.1186/s13072-021-00391-x

54. Sabirov, M., Popovich, A., Boyko, K., Nikolaeva, A., Kyrchanova, O., Maksimenko, O., Popov, V., Georgiev, P., and Bonchuk, A. 2021b. Mechanisms of CP190 Interaction with Architectural Proteins in Drosophila Melanogaster. Int J Mol Sci 22. DOl: 10.3390/ijms222212400

55. Scheres, S.H. 2012. A Bayesian view on cryo-EM structure determination. J Mol Biol 415: 406–418. DOl: 10.1016/j.jmb.2011.11.010

56. Siegmund, T., and Lehmann, M. 2002. The Drosophila Pipsqueak protein defines a new family of helix-turn-helix DNA-binding proteins. Dev Genes Evol 212: 152–157. DOl: 10.1007/s00427-002-0219-2

57. Siggs, O.M., and Beutler, B. 2012. The BTB-ZF transcription factors. Cell Cycle 11: 3358–3369. DOl: 10.4161/cc.21277

58. Silva, D., Olsen, K.W., Bednarz, M.N., Droste, A., Lenkeit, C.P., Chaharbakhshi, E., Temple-Wood, E.R., and Jemc, J.C. 2016. Regulation of Gonad Morphogenesis in Drosophila melanogaster by BTB Family Transcription Factors. PLoS One 11: e0167283. DOl: 10.1371/journal.pone.0167283

59. Stead, M.A., Trinh, C.H., Garnett, J.A., Carr, S.B., Baron, A.J., Edwards, T.A., and Wright, S.C. 2007. A beta-sheet interaction interface directs the tetramerisation of the Miz-1 POZ domain. J Mol Biol 373: 820–826. DOl:

60. Stead, M.A., and Wright, S.C. 2014. Structures of heterodimeric POZ domains of Miz1/BCL6 and Miz1/NAC1. Acta Crystallogr F Struct Biol Commun 70: 1591–1596. DOl: 10.1107/S2053230X14023449

61. Stogios, P.J., Chen, L., and Prive, G.G. 2007. Crystal structure of the BTB domain from the LRF/ZBTB7 transcriptional regulator. Protein Sci 16: 336–342. DOl: 10.1110/ps.062660907

62. Stogios, P.J., Cuesta-Seijo, J.A., Chen, L., Pomroy, N.C., and Prive, G.G. 2010. Insights into strand exchange in BTB domain dimers from the crystal structures of FAZF and Miz1. J Mol Biol 400: 983–997. DOl: 10.1016/j.jmb.2010.05.028S0022-2836(10)00507-3 [pii]

63. Stogios, P.J., Downs, G.S., Jauhal, J.J., Nandra, S.K., and Prive, G.G. 2005. Sequence and structural analysis of BTB domain proteins. Genome Biol 6: R82. DOl:

64. Svergun, D.I., Barberato, C., and Koch, M.H.J. 1995. CRYSOL - a Program to Evaluate X-ray Solution Scattering of Biological Macromolecules from Atomic Coordinates J Appl Cryst 28: 768-773. DOl:

65. Tang, X., Li, T., Liu, S., Wisniewski, J., Zheng, Q., Rong, Y., Lavis, L.D., and Wu, C. 2022. Kinetic principles underlying pioneer function of GAGA transcription factor in live cells. Nat Struct Mol Biol 29: 665–676. DOl: 10.1038/s41594-022-00800-z

66. Tikhonov, M., Utkina, M., Maksimenko, O., and Georgiev, P. 2018. Conserved sequences in the Drosophila mod(mdg4) intron promote poly(A)-independent transcription termination and trans-splicing. Nucleic Acids Res 46: 10608–10618. DOl: 10.1093/nar/gky716

67. Vogelmann, J., Le Gall, A., Dejardin, S., Allemand, F., Gamot, A., Labesse, G., Cuvier, O., Negre, N., Cohen-Gonsaud, M., Margeat, E., and Nollmann, M. 2014. Chromatin insulator factors involved in long-range DNA interactions and their role in the folding of the Drosophila genome. PLoS Genet 10: e1004544. DOl: 10.1371/journal.pgen.1004544

68. Yu, G. 2020. Using ggtree to Visualize Data on Tree-Like Structures. Curr Protoc Bioinformatics 69: e96. DOl: 10.1002/cpbi.96

69. Zacharchenko, T., and Wright, S. 2021. Functionalization of the BCL6 BTB domain into a noncovalent crystallization chaperone. IUCrJ 8: 154–160. DOl: 10.1107/S2052252520015754

70. Zollman, S., Godt, D., Prive, G.G., Couderc, J.L., and Laski, F.A. 1994. The BTB domain, found primarily in zinc finger proteins, defines an evolutionarily conserved family that includes several developmentally regulated genes in Drosophila. Proc Natl Acad Sci U S A 91: 10717–10721. DOl: 10.1073/pnas.91.22.10717

