## Supplementary Figures and Tables S1-6 for "The Arthropoda-specific Tramtrack group BTB protein domains use previously unknown interface to form hexamers"

**Supplementary Tables**

**Supplementary Table S1**. Oligonucleotides used for cloning. Restriction enzyme sites are shown in small letters, the corresponding enzymes are noted. Nucleotide substitutions in mutagenic primers are also shown in small letters.

| **Name** | **Sequence** | **restriction enzyme** |
|---|---|---|
| Bcl6 BTBdir | TAagatctAAAATGGCCTCGCCGG | BglII |
| Bcl6 BTBrev | TTAagcgctGGCCTTAATAAACTTCCG | *Eco47III* |
| GAGAdir | TAagatctACGATGTCGCTGCCAATG | *BglII* |
| GAGA 120 rev | ATggatccCGTCTGCGGTGCCAG | *BamHI* |
| batman_d | TTCTtcatgaTGTCGTCGGATC | *PagI* |
| batman_r | AATActcgagACCTTCCCGCTTTA | *Xho*I |
| psq_d | TACGccatggCAGCGGTTC | *NcoI* |
| psq_120r | CGTTctcgagGGTTTCGCAGA | *XhoI* |
| FRU_d | CAAggatccATGGACCAGCAATTCTGCTTG | *BamHI* |
| FRU_118r | GTActcgagATTGTTGTTATCTGTGAGACCAC | *XhoI* |
| LOLA_d | GAAggatccATGGATGACGATCAGCAG | *BamHI* |
| LOLA_120r | TCCctcgagAGTGCGATTGTCCGAAAGG | *XhoI* |
| BRC_d | ATCggatccATGGACGACACACAGCACTTC | *BamHI* |
| BRC_121r | ATGctcgagCTCTGCCTGCTGCTGCGTG | *XhoI* |
| BAB1_97d | ATCggatccAGCTCCCAGCAATTCTG | *BamHI* |
| BAB1_215r | AGCctcgagATGGGTCACATCCGCCAGAC | *XhoI* |
| BAB2_194d | ATCggatccGAGGGGCAGCAGTTCTGC | *BamHI* |
| BAB2_313r | TTCctcgagCCTGCCCGCACTGACCTC | *XhoI* |
| BTB VIId | CTCggatccATGTCCGTGCAGCAGTTC | *BamHI* |
| BTBVII-119r | TAActcgagttaCGTTGGCATCTCCGCTAGG | *XhoI* |
| 6765d | AACggatccATGGCCGCCGAAAACTATCAC | *BamHI* |
| 6765_133r | ATTctcgagTTAGGTGCGGCAGAGGCCAC | *XhoI* |
| ribbon_d | AAggatccATGGGCGGCCCAACGGCG | *BamHI* |
| ribbon_137r | TTAGCTGGCATGATTGAACTTCATC | |
| chinmo_d | CTagatctATGGATCCGCAGCAGCAGTTC | *BglII* |
| chinmo121r | ttaTCCCGTTTCCGTGGACAGAC | |
| 3726_d | GCggatccATGCTGCCGCAGCAGTAC | *BamHI* |
| 3726_125r | TTAGTCTTCGTCATCGCGCCAGG | |
| mamo_d | ATCggatccATGGGCAGTGAGCACTACTG | *BamHI* |
| mamo_121r | AGctcgagTTACTGGTTCGTCATCTCGGCCAG | *XhoI* |
| 6118_306d | TTCggatccATCGATCAGTACCTGC | *BamHI* |
| 6118_419r | CAGgtcgacCTATGTGTGCAGCCCCTTG | *SalI* |
| CG15812d | CACggatccATGAATCATCTTAAGTGGATG | *BamHI* |
| CG15812_Br | TGGgtcgacTTAAAAGCTGATGGCAGATTTG | *SalI* |
| CG34376d | TCAggatccATGGACGACGAGTTTAAGC | *BamHI* |
| CG34376_Br | CGAgtcgacTTAATTGGTAGCCAAGCCCTTG | *SalI* |
| TTK_d | GTCggatccATGAAGATGGCATCTCAACG | *BamHI* |
| TTK_117r | ATCgtcgacGGTGAGGCCCTTGATGCG | *SalI* |
| CG15275d | attggatccATGTTCACCAATTGGCTGACTG | *BamHI* |
| CG15275_191r | atactcgagttaTTGAGCGCGCGCCAAAAATGAGCG | *XhoI* |
| CG6792d | GCTGgatccATGCTGCACTCACAGACAATGC | *BamHI* |
| CG6792_134r | CTTgtcgacttaCAAGGGCTGCAACTGGTAGAG | *SalI* |
| Ken1d | TCCggatccATGAAAGAGTTTCAAAGAATGTTG | *BamHI* |
| Ken147r | tatgtcgacttaGTACTGCTGCTTCCGCCTG | *SalI* |
| 6765_Y84K V86Kd | GCTGaagGTGaagCTGCCGCCGGATC | |

| | |
|---|---|
| 6765_Y84K V86Kr | GCAGcttCACcttCAGCACTCCATTGGGAT |
| LOLA I70 I72Ad | CCCgccTTTgcaCTCAAGGATGTCAAGTAC |
| LOLA I70 I72Ar | CTTGAGtgcAAAggcGGGATGTTTGTCGTACTG |
| LOLA I70 I72Kd | CCCaagTTTaaaCTCAAGGATGTCAAGTAC |
| LOLA I70 I72Kr | CCTTGAGtttAAActtGGGATGTTTGTCGTACTG |
| MOD I71K F73Kd | CGCTaagGTAaagCTGAACAACGTCAGCCAC |
| MOD I71K F73Kr | GTTGTTCAGcttTACcttAGCGTGGGTGTTCGAC |
| MOD I71A F73Ad | GCTgccGTAgccCTGAACAACGTCAGCC |
| MOD I71A F73Ar | CAGggcTACggcAGCGTGGGTGTTCGAC |
| MOD_F73Ad | GCTATCGTAgcCCTGAACAACGTCAGCC |
| MOD_F73Ar | CGTTGTTCAGGgcTACGATAGCGTGGGTG |
| MOD_F73Pd | GCTATCGTAccCCTGAACAACGTCAGCC |
| MOD_F73Pr | CGTTGTTCAGGggTACGATAGCGTGGGTG |
| LOLA72Ad | CCCATCTTTgcACTCAAGGATGTCAAGTAC |
| LOLA72Ar | CTTGAGTgcAAAGATGGGATGTTTGTCGTACTG |
| LOL70P72Ad | CCCccCTTTgcACTCAAGGATGTCAAGTAC |
| LOL70P72Ar | CTTGAGTgcAAAGggGGGATGTTTGTCGTACTG |

**Supplementary Table S2. Cryo-EM statistics. Data collection, processing, model refinement and validation statistics.**

| | |
|---|---|
| **Data collection and processing** | |
| Magnification | 165,000 |
| Voltage(kV) | 300 |
| Electron exposure($e^-$/Å²) | 60 |
| Energy filter slit width(eV) | — |
| Defocus range(μm) | -0.8 to -2.2 |
| Pixel size(Å) | 0.729 |
| Symmetry imposed | C1 |
| Movies(no.) | 9,765 |
| Initial particle images(no.) | 480,721 |
| Final particle images(no.) | 197,562 |
| Map resolution(Å)<br>FSC threshold. | 3.3<br>0.143 |
| **Refinement** | |
| Initial model used | AlphaFold |
| Model composition<br>Non-hydrogen atoms | <br>6118 |
| Protein residues | 780 |
| R.m.s deviations | |
| Bond lengths(Å) | 0.005 |
| Bond angles(°) | 0.553 |
| **Validation** | |
| MolProbity score | 1.80 |
| Clashscore | 6.07 |
| Rotamer outliers(%) | 0.58 |
| Ramachandran plot | |
| Favored(%) | 98.57 |
| Allowed(%) | 1.30 |
| Disallowed(%) | 0.13 |
| Model to map CC (mask, peaks, volume) | 0.77/0.76/0.66 |

**Supplementary Table S3**. The ability of GAF, Mod(mdg4), Chinmo, CG8924 and LOLA BTB domains to interact with other TTK-type BTB domains in yeast two-hybrid assay. BCL6 is heterologous BTB domain of human BCL6 protein. GAF was tested both as AD and BD fusions, whereas Mod(mdg4), Chinmo, CG8924 and LOLA were tested only as BD fusions. Interactions with CG32121 were tested only for AD fusions due to strong self-activation properties. AD stands for Activation Domain, BD – for DNA-Binding Domain of GAL4 protein. + or - denotes the ability of yeasts to grow on the media without histidine. Yeast assay plates are shown in Supplementary Figure S2.

| | | AD | | |
|---|---|---|---|---|
| | | GAF BTB | - | dimerization control |
| BD | GAF | + | – | + |
| | Abrupt | + | – | + |
| | CG3726 | + | – | + |
| | CG12236 | + | – | + |
| | BTB VII | + | – | + |
| | bab2 | + | – | + |
| | bab1 | + | – | + |
| | Ribbon | + | – | + |
| | mod | + | – | + |
| | lola | + | – | + |
| | ttk | + | – | + |
| | Psq | + | – | + |
| | BCL6 | – | – | + |
| | Batman | + | – | + |
| | CG6118 | + | – | + |
| | CG15812 | – | – | + |
| | CG34376 | + | – | + |
| | Fruitless | – | – | + |
| | TKR | + | – | + |
| | CG8924 | + | – | + |
| | mamo | + | – | + |
| | BRC | + | – | + |
| | Chinmo | – | – | + |
| | CG6765 | + | – | + |
| | - | – | – | |

| | | BD | | | | | |
|---|---|---|---|---|---|---|---|
| | | mod(mdg4) BTB | LOLA BTB | GAF BTB | CG8924 BTB | Chinmo BTB | - |
| AD | mod(mdg4) | + | – | + | – | – | – |
| | CG32121 | + * | + | + | – | – | – |
| | Abrupt | – | – | + | – | + | – |
| | CG3726 | + | + | + | + | + | – |
| | CG12236 | – | – | – | – | + | – |
| | BTB VII | + | + | + | – | – | – |
| | bab2 | + | – | + | + | + | – |
| | bab1 | + | – | + | + | + | – |
| | Ribbon | + | + | – | + | + | – |
| | GAF | + | + | + | – | – | – |
| | lola | – | + | + | – | – | – |
| | ttk | – | + | + | + | + | – |
| | Psq | – | – | + | + | + | – |
| | BCL6 | – | – | – | – | – | – |
| | Batman | + | – | + | + | + | – |
| | CG6118 | + | – | + | + | + | – |
| | CG15812 | + | – | + | + | + | – |
| | CG34376 | + | – | + | + | – | – |
| | Fruitless | + | + | – | + | + | – |
| | TKR | + | + | + | + | + | – |
| | CG8924 | – | – | + | + | + | – |
| | mamo | – | + | + | + | + | – |
| | BRC | + | – | + | – | + | – |
| | Chinmo | – | – | – | – | + | – |
| | CG6765 | – | – | + | – | + | – |
| | - | – | – | – | – | – | – |

**Supplementary Table S4.** Pairwise level of similarity between GAF and Mod(mdg4) BTB domains with other TTK-type domains. The level of homology was calculated as ratio of identical and conserved (according to CLUSTAL definitions) residues to the total number of residues.

| | GAF | mod(mdg4) |
|---|---|---|
| GAF | 100 | 35 |
| CG32121 | 36 | 39 |
| Abrupt | 47 | 40 |
| CG3726 | 35 | 40 |
| CG12236 | 35 | 42 |
| BTB VII | 40 | 47 |
| bab2 | 37 | 43 |
| bab1 | 42 | 42 |
| Ribbon | 34 | 35 |
| mod (mdg4) | 35 | 100 |
| lola | 38 | 45 |
| ttk | 38 | 47 |
| Psq | 38 | 37 |
| Batman | 39 | 44 |
| CG6118 | 33 | 47 |
| CG15812 | 27 | 25 |
| CG34376 | 39 | 45 |
| Fruitless | 31 | 51 |
| TKR | 37 | 36 |
| CG8924 | 33 | 37 |
| mamo | 38 | 44 |
| BRC | 46 | 43 |
| Chinmo | 32 | 40 |
| CG6765 | 27 | 32 |

**Supplementary Table S5.** Testing of the interactions between the non-TTK BTB domains of C2H2 proteins. (*) – the growth was assessed in the presence of 3-aminotriazole due to strong self-activation properties. Yeast assay plates are shown in Supplementary Figure S2.

| | | AD | | | | |
|---|---|---|---|---|---|---|
| | | CP190 | ken | CG6792 | CG15275 | - |
| BD | CP190 | **+** | – | – | – | – |
| | ken | **-** | **+** | **-** | – | – |
| | CG6792* | – | – | **+** | – | – |
| | CG15275 * | – | – | – | **+** | – |
| | - | – | – | – | – | – |

**Supplementary Table S6.** The ability of non-ttk BTB domains of CP190, CG6792, CG15725 and Ken proteins to interact with other TTK-type BTB domains in yeast two-hybrid assay. Positive results from initial screen against AD-tagged baits were further studied in reciprocal experiment as BD-tagged (lower table). BCL6 is heterologous BTB domain of human BCL6 protein used as an additional control. Designations are the same as in Supplementary Table S2. Yeast assay plates are shown in Supplementary Figure S4.

| | | AD | | | | |
|---|---|---|---|---|---|---|
| | | CP190 BTB | Ken BTB | CG6792 BTB | CG15725 BTB | - |
| BD | mod(mdg4) | – | – | – | – | – |
| | CG32121 | –* | –* | –* | –* | –* |
| | Abrupt | – | – | – | – | – |
| | CG3726 | – | – | – | – | – |
| | CG12236 | – | – | – | – | – |
| | BTB VII | – | – | – | – | – |
| | bab2 | – | – | – | – | – |
| | bab1 | – | – | – | – | – |
| | Ribbon | + | – | – | – | – |
| | GAF | + | – | – | – | – |
| | lola | – | – | – | – | – |
| | ttk | – | – | – | – | – |
| | Psq | – | – | – | – | – |
| | BCL6 | – | – | – | – | – |
| | Batman | – | + | – | – | – |
| | CG6118 | – | – | – | – | – |
| | CG15812 | – | – | – | – | – |
| | CG34376 | – | – | – | – | – |
| | Fruitless | – | – | – | – | – |
| | TKR | – | – | – | – | – |
| | CG8924 | – | – | – | – | – |
| | mamo | – | + | – | – | – |
| | BRC | – | – | – | – | – |
| | Chinmo | – | – | – | – | – |
| | CG6765 | – | – | – | – | – |
| | - | – | – | – | – | – |

| | | BD | |
|---|---|---|---|
| | | CP190 BTB | - |
| AD | Ribbon | + | – |
| | GAF | – | – |
| | - | – | – |

### Supplementary Figures

**Supplementary Figure S1. Flowchart illustrating cryo-EM data processing of MBP-fused CG6765$^{1-133}$. Details are described in the Methods section.**

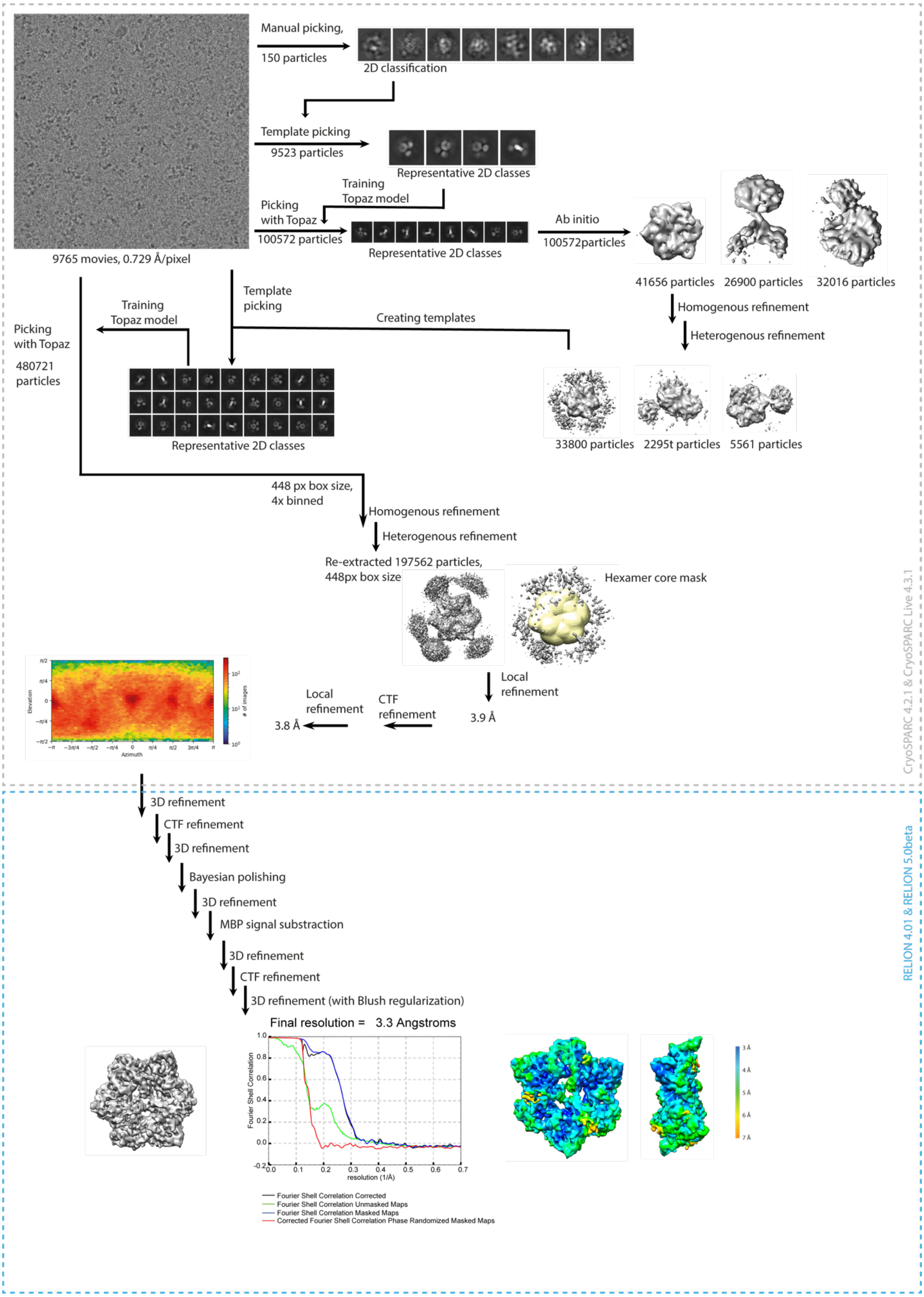

**Supplementary Figure S2. Results of yeast two-hybrid assays.** Growth assay plates without histidine are shown (yeasts are unable to grow on this medium in the absence of interaction). AD stands for Activation Domain, BD – for DNA-Binding Domain of GAL4 protein. Unlabeled plate segments correspond to excessive controls.

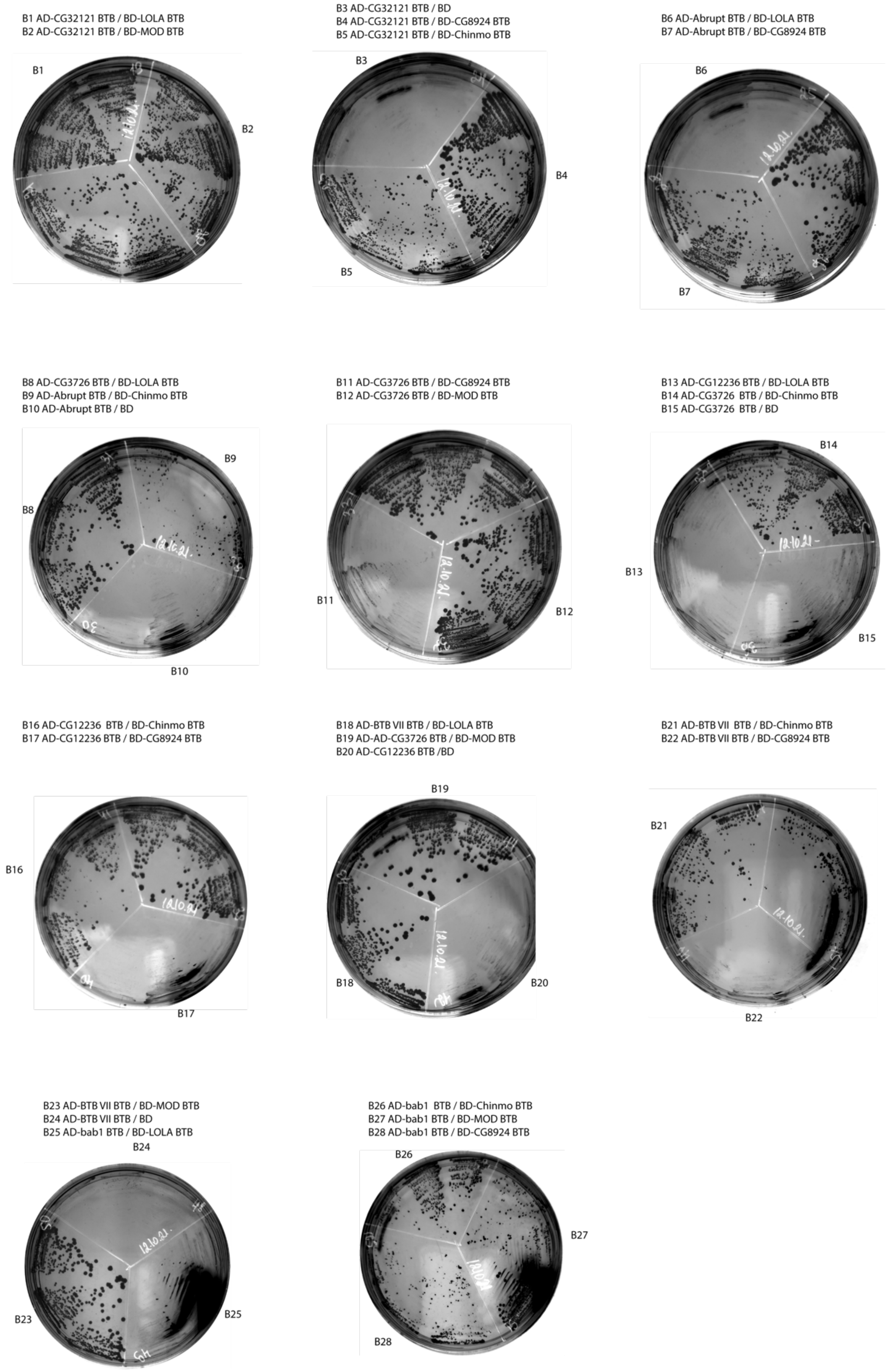

A1 AD-bab1 BTB / BD
A2 AD-bab2 BTB / BD-LOLA BTB
A3 AD-bab2 BTB / BD-MOD BTB

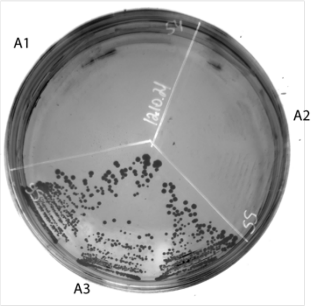

A4 AD-bab2 BTB / BD-CG8924 BTB
A5 AD-bab2 BTB / BD-Chinmo BTB

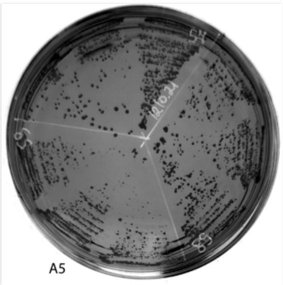

A6 AD-bab1 BTB / BD
A7 AD-Ribbon BTB / BD-LOLA BTB
A8 AD-Ribbon BTB / BD-MOD BTB

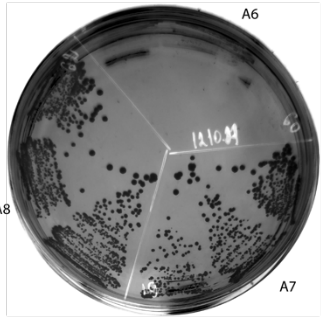

A9 AD-Ribbon BTB / BD-CG8924 BTB
A10 AD-Ribbon BTB / BD-Chinmo BTB

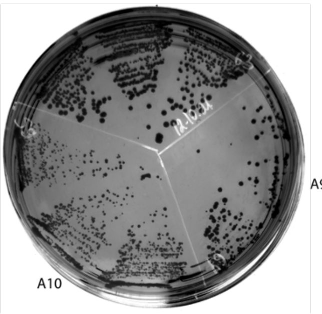

A11 AD-Ribbon BTB / BD
A12 AD-Fruitless BTB / BD-LOLA BTB
A13 AD-Fruitless BTB / BD-MOD BTB

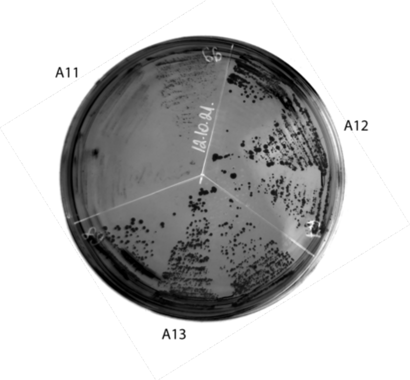

A14 AD-Fruitless BTB / BD-CG8924 BTB
A15 AD-Fruitless BTB / BD-Chinmo BTB

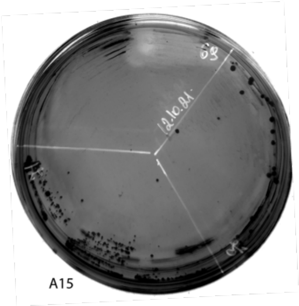

A16 AD-Fruitless BTB / BD
A17 AD-TKR BTB / BD-LOLA BTB
A18 AD-TKR BTB / BD-MOD BTB

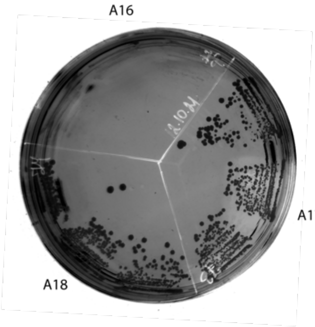

A19 AD-TKR BTB / BD-CG8924 BTB
A20 AD-TKR BTB / BD-Chinmo BTB

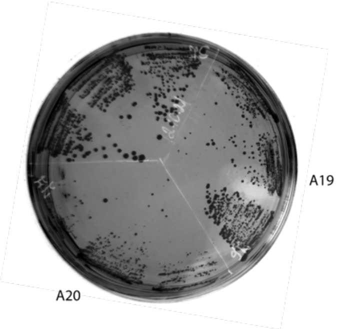

A21 AD-TKR BTB / BD
A22 AD-CG8924 BTB / BD-LOLA BTB

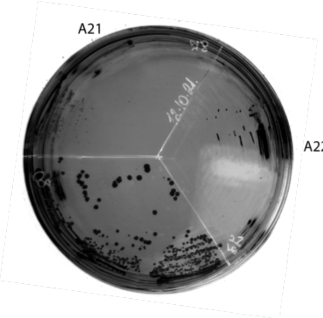

A23 AD-CG8924 BTB / BD-CG8924 BTB
A24 AD-CG8924 BTB / BD-Chinmo BTB
A25 AD-CG8924 BTB / BD

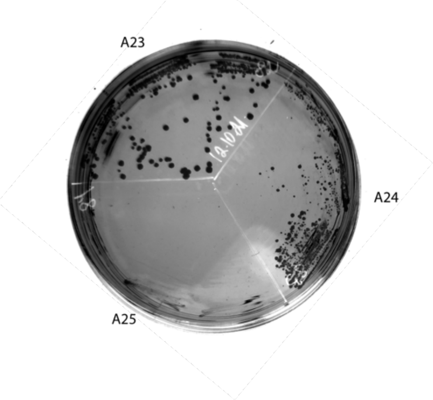

A26 AD-MAMO BTB / BD-LOLA BTB
A27 AD-MAMO BTB / BD-CG8924 BTB

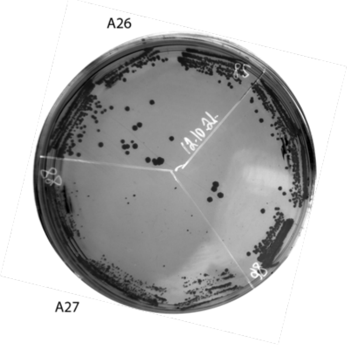

A28 AD-BRC BTB / BD-LOLA BTB
A29 AD-MAMO BTB / BD-Chinmo BTB
A30 AD-MAMO BTB / BD

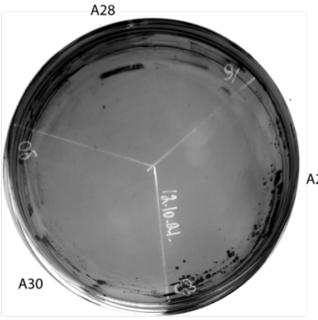

C1 AD-BRC BTB / BD-MOD BTB
C2 AD-BRC BTB / BD-CG8924 BTB

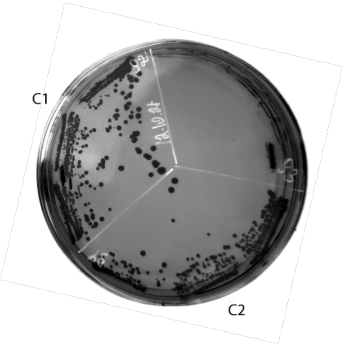

C3 AD-BRC BTB / BD
C4 AD-Chinmo BTB / BD-LOLA BTB
C5 AD-BRC BTB / BD-Chinmo BTB

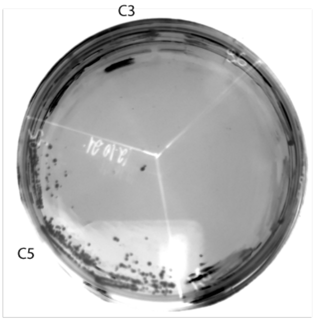

C6 AD-Chinmo BTB / BD-CG8924 BTB
C7 AD-Chinmo BTB / BD-Chinmo BTB

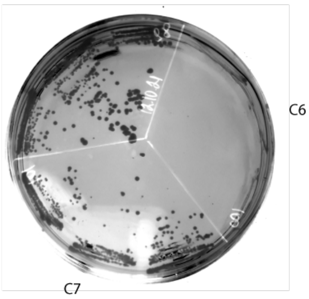

C8 AD-Chinmo BTB / BD
C9 AD-CG6765 BTB / BD-LOLA BTB

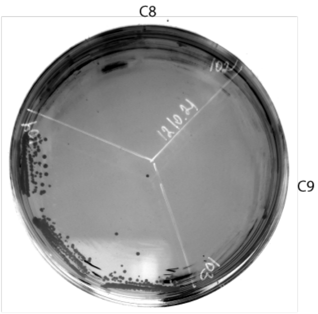

C10 AD-CG6765 BTB / BD-CG8924 BTB
C11 AD-CG6765 BTB / BD-Chinmo BTB
C12 AD-CG6765 BTB / BD

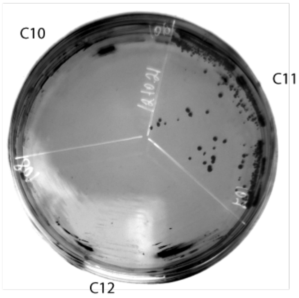

C13 AD-CG15812 BTB / BD-LOLA BTB
C14 AD-CG15812 BTB / BD-MOD BTB

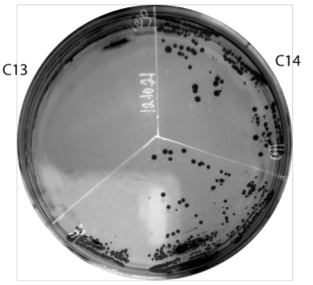

C15 AD-CG15812 BTB / BD-CG8924 BTB
C16 AD-CG15812 BTB / BD-Chinmo BTB
C17AD-CG15812 BTB / BD

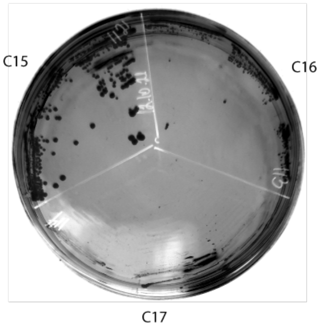

C18 AD-CG6118 BTB / BD-LOLA BTB
C19 AD-CG6118 BTB / BD-MOD BTB

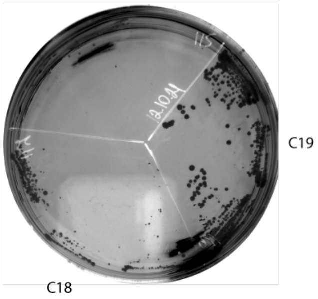

C20 AD-CG6118 BTB / BD-CG8924 BTB
C21 AD-CG6118 BTB / BD-Chinmo BTB
C22 AD-CG6118 BTB / BD

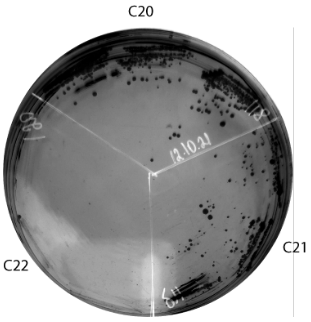

C23 BD-GAF BTB / AD-GAF BTB
C24 BD-GAF BTB / AD
C25 BD-GAF BTB / AD-CG32121 BTB
C26 BD-Abrupt BTB / AD-GAF BTB

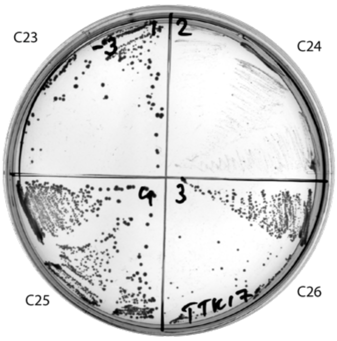

C27 BD-Abrupt BTB / AD
C28 BD-CG3726 BTB / AD-GAF BTB
C29 BD-CG3726 BTB / AD
C30 BD-CG12236 BTB / AD-GAF BTB

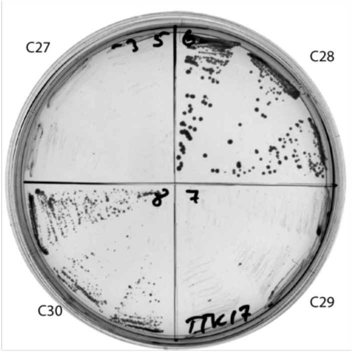

C31 BD-CG12236 BTB / AD
C32 BD-BTB VII BTB / AD-GAF BTB
C33 BD-BTB VII BTB / AD
C34 BD-bab2 BTB / AD-GAF BTB

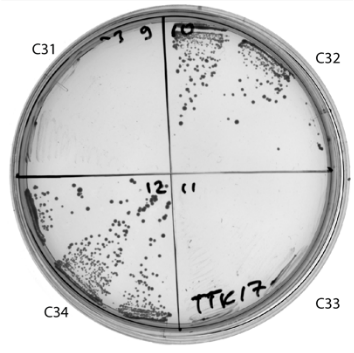

C35 BD-bab2 BTB / AD
C36 BD-bab1 BTB / AD-GAF BTB
C37 BD-bab1 BTB / AD
C38 BD-Ribbon BTB / AD-GAF BTB

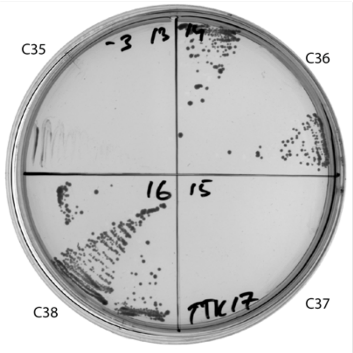

C39 BD-Ribbon BTB / AD
C40 BD-Mod(mdg4) BTB / AD-GAF BTB
C41 BD-Mod(mdg4) BTB / AD
C42 BD-lola BTB / AD-GAF BTB

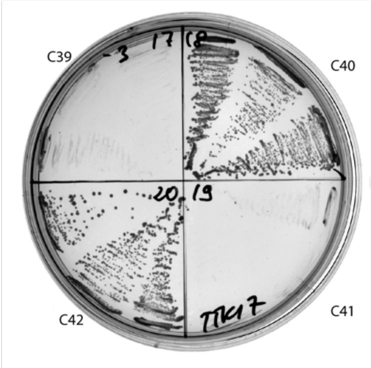

C43 BD-lola BTB / AD
C44 BD-Tramtrack BTB / AD-GAF BTB
C45 BD-Tramtrack / AD
C46 BD-Pipsqueak BTB / AD-GAF BTB

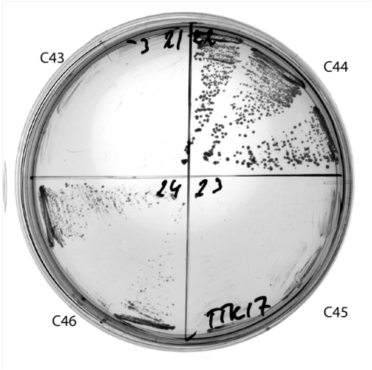

C47 BD-Pipsqueak BTB / AD
C48 BD-BCL6 BTB / AD-GAF BTB
C49 BD-BCL6 BTB / AD
C50 BD-Batman BTB / AD-GAF BTB

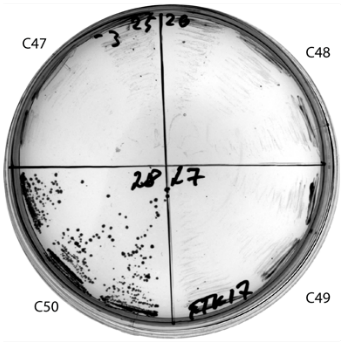

C51 BD-Batman BTB / AD
C52 BD-CG6118 BTB / AD-GAF BTB
C53 BD-CG6118 BTB / AD
C54 BD-CG15812 BTB / AD-GAF BTB

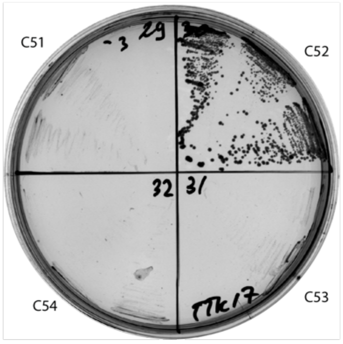

C55 BD-CG15812 BTB / AD
C56 BD-CG34376 BTB / AD-GAF BTB
C57 BD-CG34376 / AD
C58 BD-Fruitless BTB / AD-GAF BTB

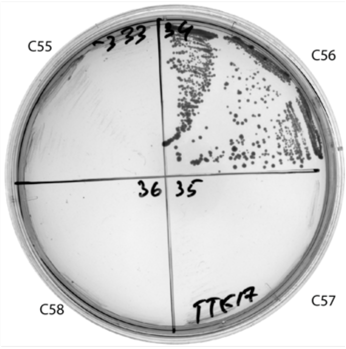

C59 BD-Fruitless BTB / AD
C60 BD-TKR BTB / AD-GAF BTB
C61 BD-TKR BTB / AD
C62 BD-CG8924 BTB / AD-GAF BTB

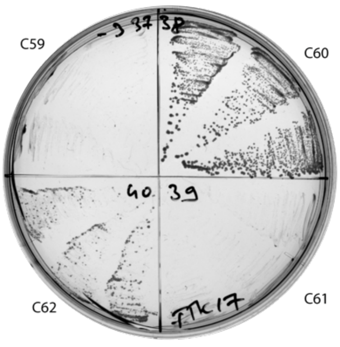

C63 BD-CG8924 BTB / AD
C64 BD-mamo BTB / AD-GAF BTB
C65 BD-mamo BTB / AD
C66 BD-BR-C BTB / AD-GAF BTB

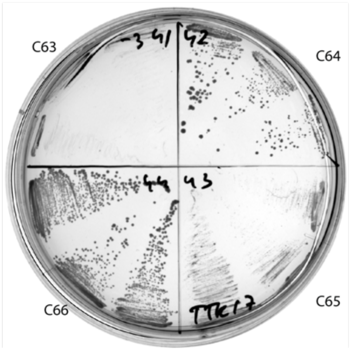

C67 BD-BR-C BTB / AD
C68 BD-Chinmo BTB / AD-GAF BTB
C69 BD-Chinmo BTB / AD
C70 BD-CG6765 BTB / AD-GAF BTB

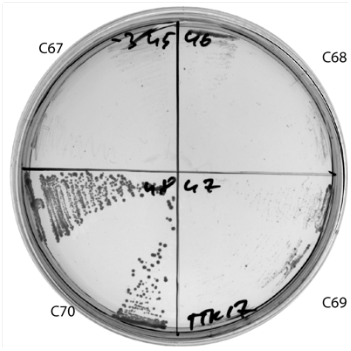

C71 BD-CG6765 BTB / AD
C72 BD / AD-GAF BTB
C73 BD / AD
C74 BD / AD-CG32121 BTB

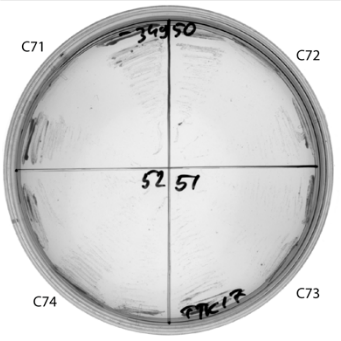

C75 BD-GAF BTB / AD-GAF BTB
C76 BD / AD-GAF BTB
C77 BD-GAF BTB / AD-Abrupt BTB
C78 BD / AD-Abrupt BTB

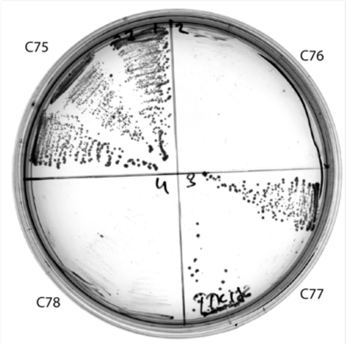

C79 BD-GAF BTB / AD-CG3726 BTB
C80 BD / AD-CG3726 BTB
C81 BD-GAF BTB / AD-CG12236 BTB
C82 BD / AD-CG12236 BTB

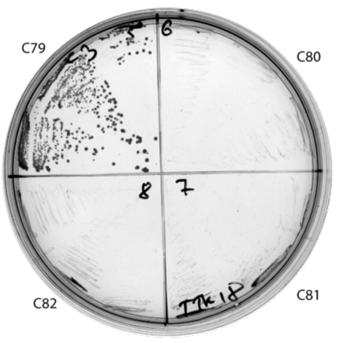

C83 BD-GAF BTB / AD-BTB VII BTB
C84 BD / AD-BTB VII BTB
C85 BD-GAF BTB / AD-bab2 BTB
C86 BD / AD-bab2 BTB

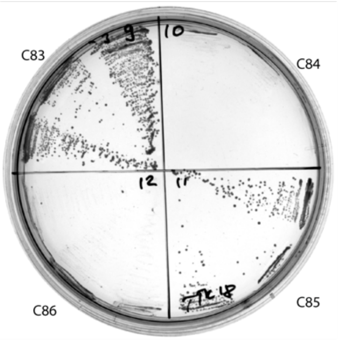

C87 BD-GAF BTB / AD-bab1 BTB
C88 BD / AD-bab1 BTB
C89 BD-GAF BTB / AD-Ribbon BTB
C90 BD / AD-Ribbon BTB

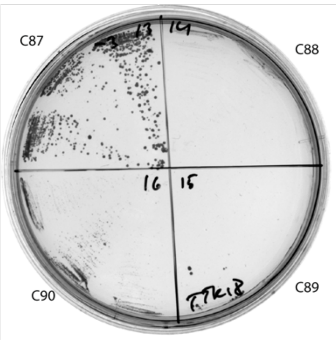

C91 BD-GAF BTB / AD-Mod(mdg4) BTB
C92 BD / AD-Mod(mdg4) BTB
C93 BD-GAF BTB / AD-lola BTB
C94 BD / AD-lola BTB

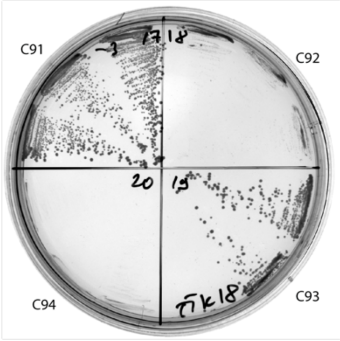

C95 BD-GAF BTB / AD-Tramtrack BTB
C96 BD / AD-Tramtrack BTB
C97 BD-GAF BTB / AD-Pipsqueak BTB
C98 BD / AD-Pipsqueak BTB

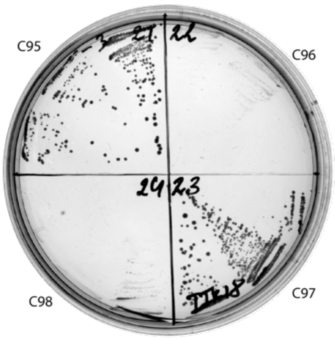

C99 BD-GAF BTB / AD-BCL6 BTB
C100 BD / AD-BCL6 BTB
C101 BD-GAF BTB / AD-Batman BTB
C102 BD / AD-Batman BTB

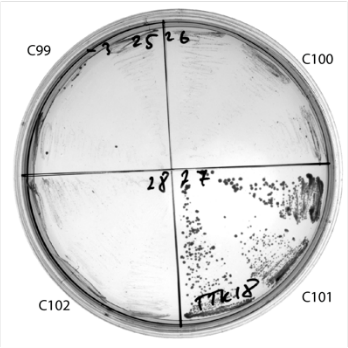

C103 BD-GAF BTB / AD-CG6118 BTB
C104 BD / AD-CG6118 BTB
C105 BD-GAF BTB / AD-CG15812 BTB
C106 BD / AD-CG15812 BTB

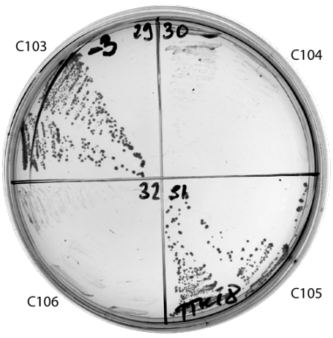

C107 BD-GAF BTB / AD-CG34376 BTB
C108 BD / AD-CG34376 BTB
C109 BD-GAF BTB / AD-Fruitless BTB
C110 BD / AD-Fruitless BTB

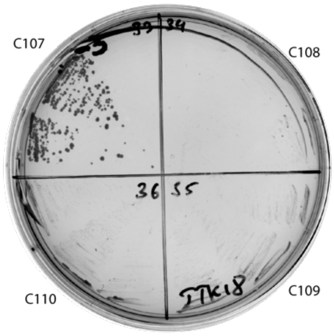

C111 BD-GAF BTB / AD-TKR BTB
C112 BD / AD-TKR BTB
C113 BD-GAF BTB / AD-CG8924 BTB
C114 BD / AD-CG8924 BTB

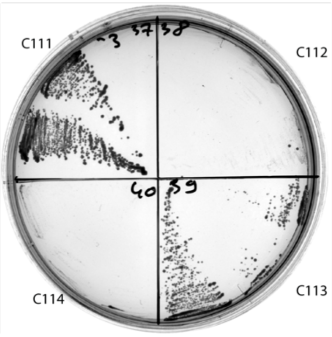

C115 BD-GAF BTB / AD-mamo BTB
C116 BD / AD-mamo BTB
C117 BD-GAF BTB / AD-BR-C BTB
C118 BD / AD-BR-C BTB

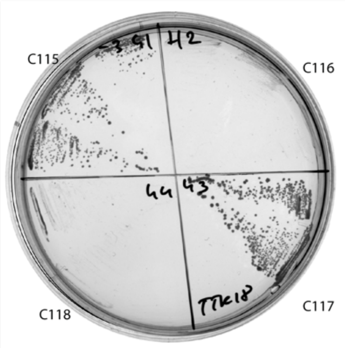

C119 BD-GAF BTB / AD-Chinmo BTB
C120 BD / AD-Chinmo BTB
C121 BD-GAF BTB / AD-CG6765 BTB
C122 BD / AD-CG6765 BTB

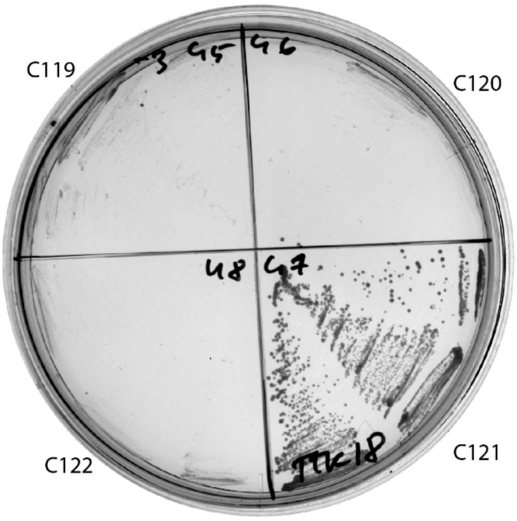

C123 BD-GAF BTB / AD
C124 BD / AD

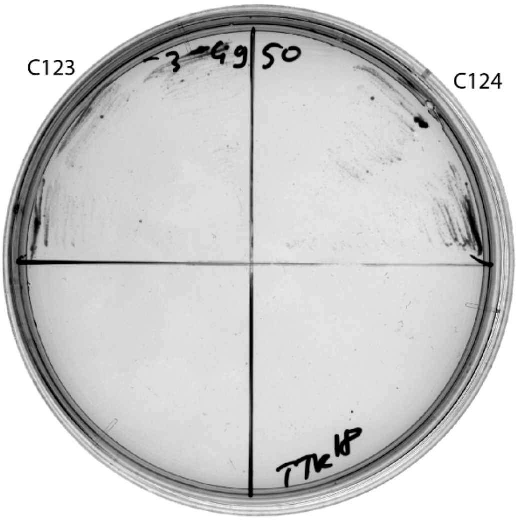

**Supplementary Figure S3. Superdex S200 size-exclusion chromatography of the CP190 and CG6792 BTB domains.** Positions of molecular weight markers are shown below. Molecular weights of monomers are shown in brackets.

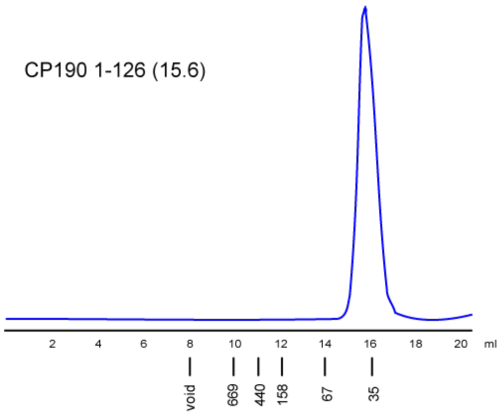

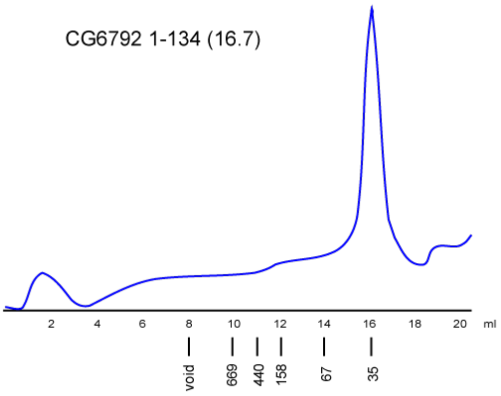

**Supplementary Figure S4. Results of yeast two-hybrid assays of interactions between TTK and non-TTK BTB domains.** Growth assay plates without histidine are shown (yeasts are unable to grow on this medium in the absence of interaction). AD stands for Activation Domain, BD – for DNA-Binding Domain of GAL4 protein.

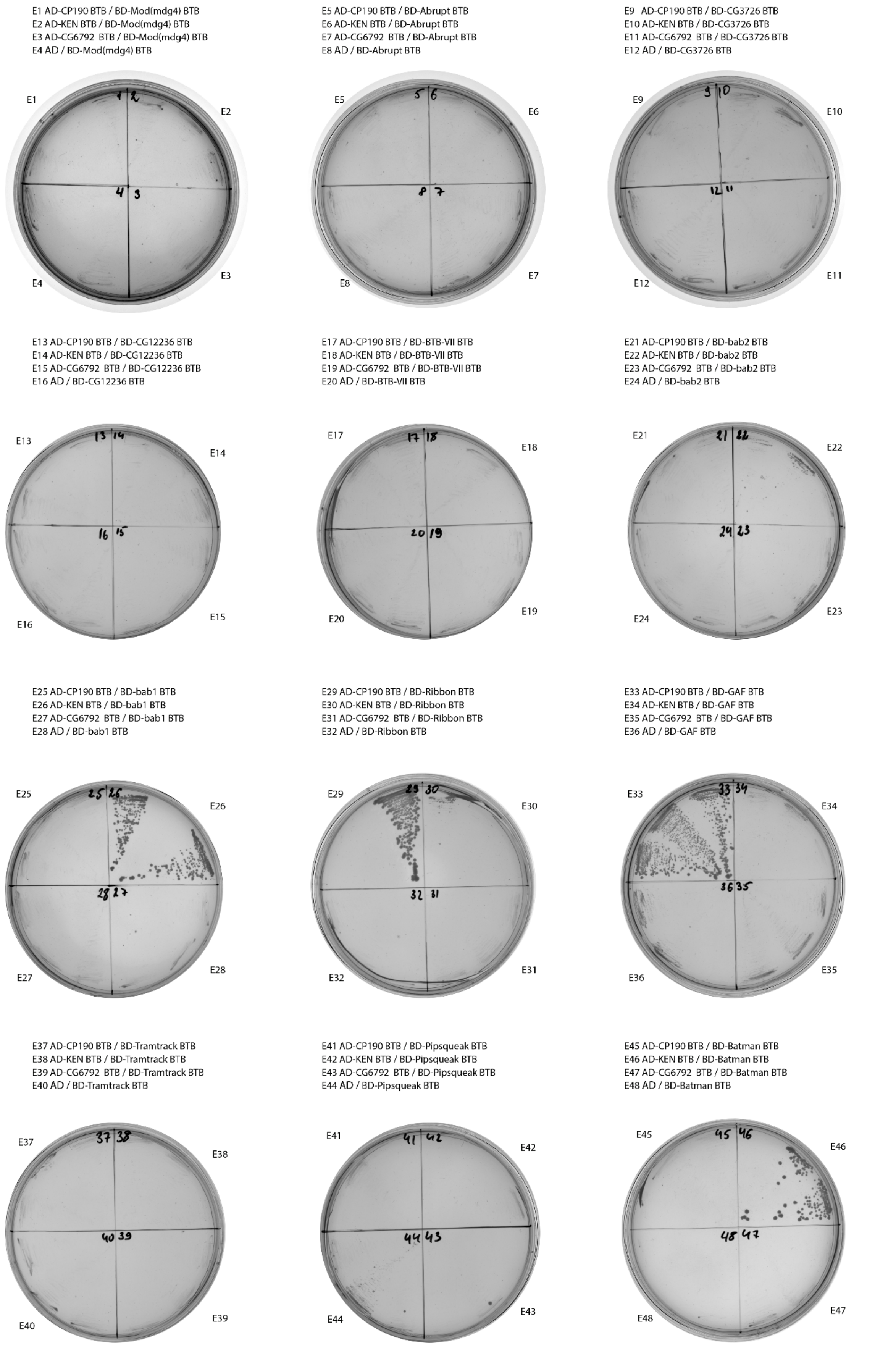

E49 AD-CP190 BTB / BD-CG6118 BTB
E50 AD-KEN BTB / BD-CG6118 BTB
E51 AD-CG6792 BTB / BD-CG6118 BTB
E52 AD / BD-CG6118 BTB

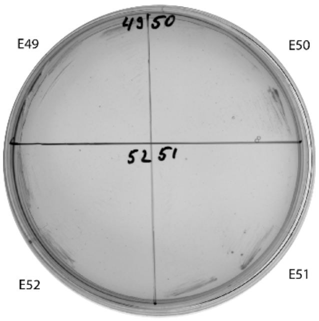

E53 AD-CP190 BTB / BD-CG15812 BTB
E54 AD-KEN BTB / BD-CG15812 BTB
E55 AD-CG6792 BTB / BD-CG15812 BTB
E56 AD / BD-CG15812 BTB

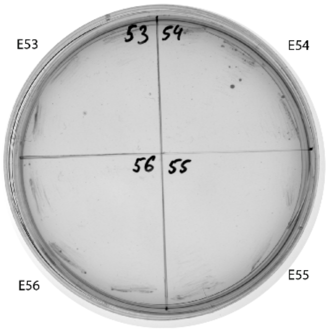

E57 AD-CP190 BTB / BD-CG34376 BTB
E58 AD-KEN BTB / BD-CG34376 BTB
E59 AD-CG6792 BTB / BD-CG34376 BTB
E60 AD / BD-CG34376 BTB

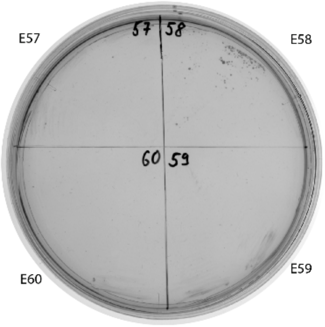

E61 AD-CP190 BTB / BD-Fruitless BTB
E62 AD-KEN BTB / BD-Fruitless BTB
E63 AD-CG6792 BTB / BD-Fruitless BTB
E64 AD / BD-Fruitless BTB

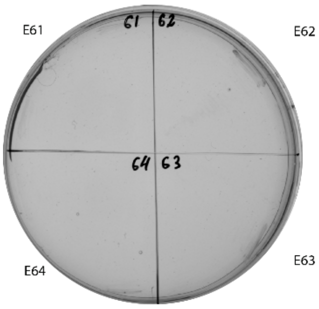

E65 AD-CP190 BTB / BD-TKR BTB
E66 AD-KEN BTB / BD-TKR BTB
E67 AD-CG6792 BTB / BD-TKR BTB
E68 AD / BD-TKR BTB

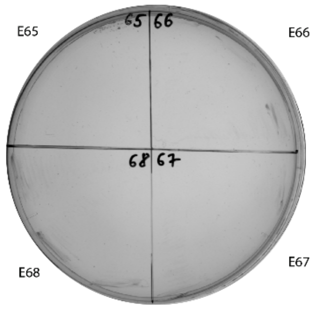

E69 AD-CP190 BTB / BD-CG8924 BTB
E70 AD-KEN BTB / BD-CG8924 BTB
E71 AD-CG6792 BTB / BD-CG8924 BTB
E72 AD / BD-CG8924 BTB

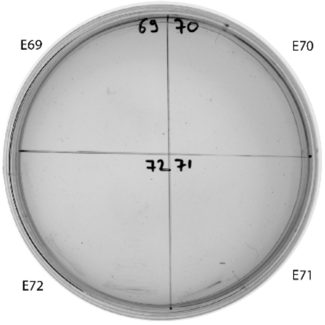

E73 AD-CP190 BTB / BD-MAMO BTB
E74 AD-KEN BTB / BD-MAMO BTB
E75 AD-CG6792 BTB /BD-MAMO BTB
E76 AD / BD-MAMO BTB

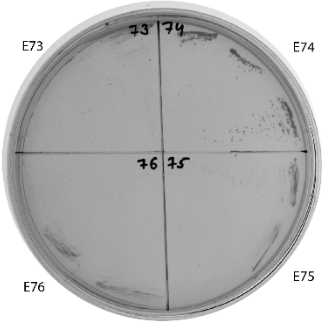

E77 AD-CP190 BTB / BD-Broad Comlex BTB
E78 AD-KEN BTB / BD-Broad Comlex BTB
E79 AD-CG6792 BTB / BD-Broad Comlex BTB
E80 AD / BD-Broad Comlex BTB

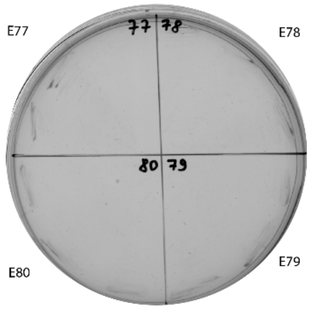

E81 AD-CP190 BTB / BD-Chinmo BTB
E82 AD-KEN BTB / BD-Chinmo BTB
E83 AD-CG6792 BTB / BD-Chinmo BTB
E84 AD / BD-Chinmo BTB

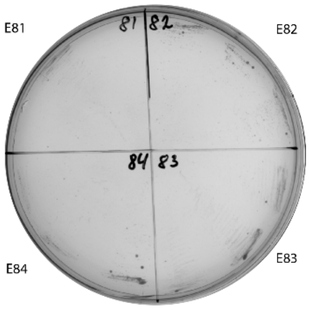

E85 AD-CP190 BTB / BD-CG6765 BTB
E86 AD-KEN BTB / BD-CG6765 BTB
E87 AD-CG6792 BTB / BD-CG6765 BTB
E88 AD / BD-CG6765 BTB

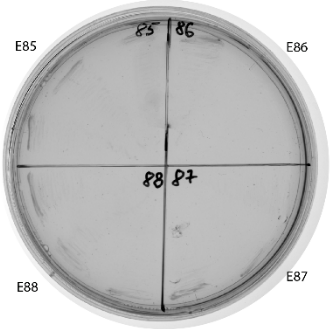

E89 AD-CP190 BTB / BD-CP190 BTB
E90 AD / BD-CP190 BTB
E91 AD-CG32121 BTB / BD-CP190 BTB
E92 AD-BCL6 BTB / BD-CP190 BTB

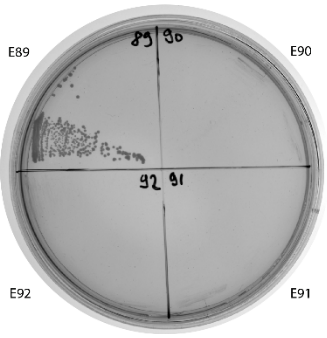

E93 AD-KEN BTB / BD-KEN BTB
E94 AD / BD-KEN BTB
E95 AD-CG32121 BTB / BD-KEN BTB
E96 AD-BCL6 BTB / BD-KEN BTB

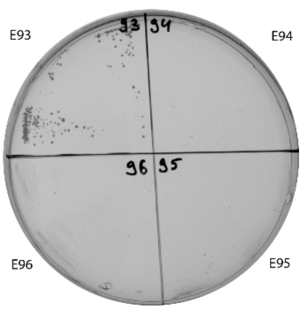

E97 AD-CG6792 BTB / BD-CG6792 BTB
E98 AD / BD-CG6792 BTB
E99 AD-CG32121 BTB / BD-CG6792 BTB
E100 AD-BCL6 BTB / BD-CG6792 BTB

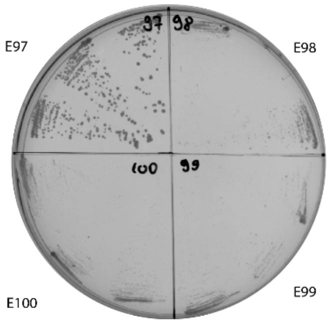

E101 AD-CP190 BTB / BD
E102 AD-KEN BTB / BD
E103 AD-CG6792 BTB / BD
E104 AD / BD

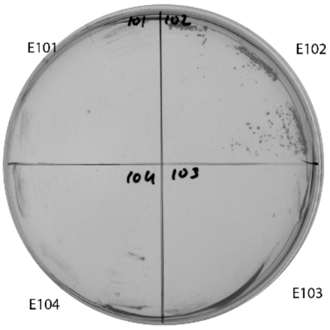

E105 AD-CG32121 BTB / BD
E106 AD-BCL6 BTB / BD

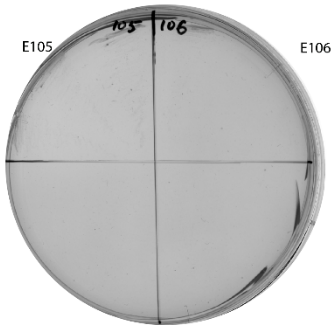

E107 AD-CP190 BTB / BD-CP190 BTB
E108 AD-Ribbon BTB / BD-CP190 BTB
E109 AD-GAF BTB / BD-CP190 BTB
E110 AD / BD-CP190 BTB

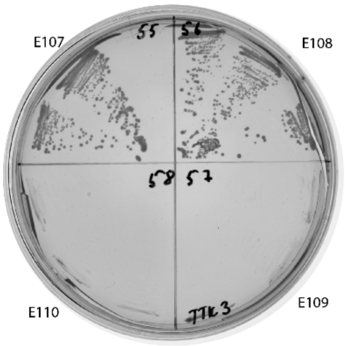

E111 AD-CP190 BTB / BD
E112 AD-Ribbon BTB / BD
E113 AD-GAF BTB / BD
E114 AD / BD

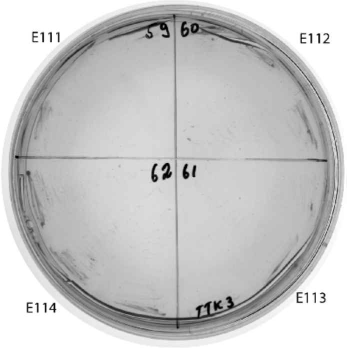

E115 BD-Ribbon BTB / AD-CP190 BTB
E116 BD-Ribbon BTB / AD
E117 BD- GAF BTB / AD-CP190 BTB
E118 BD- GAF BTB / AD

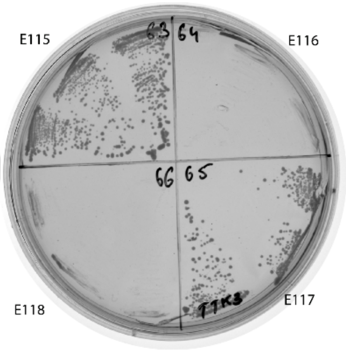

E119 BD-KEN BTB / AD-KEN BTB
E120 BD-KEN-BTB / AD
E121 BD-bab1 BTB / AD-KEN BTB
E122 BD-bab1 BTB / AD

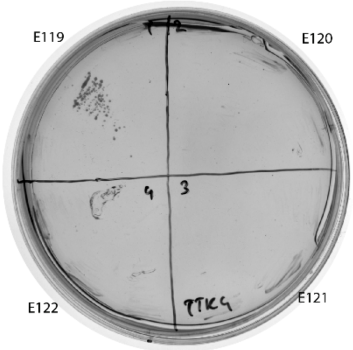

E123 BD-Batman BTB / AD-KEN BTB
E124 BD-Batman-BTB / AD
E125 BD-CG34376 BTB / AD-KEN BTB
E126 BD-CG34376 BTB / AD

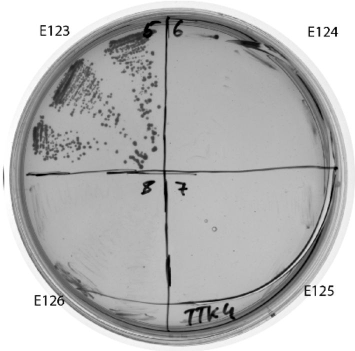

E127 BD-MAMO BTB / AD-KEN BTB
E128 BD-MAMO-BTB / AD
E129 BD-CP190 BTB / AD-CP190 BTB
E130 BD-CP190 BTB / AD

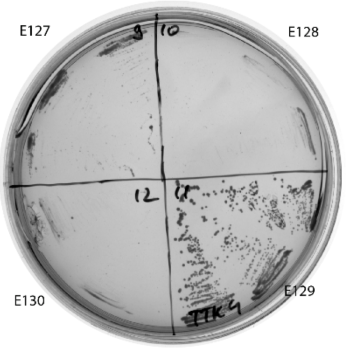

E131 BD / AD-KEN BTB
E132 BD / AD-CP190 BTB
E133 BD / AD

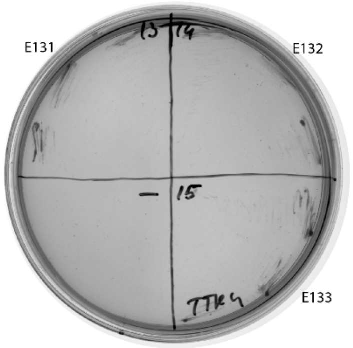

assayed in the presence of 5mM 3-aminotriazole

E134 BD-MAMO BTB / AD-KEN BTB
E135 BD-Chinmo BTB / AD-KEN BTB
E136 BD / AD-KEN BTB

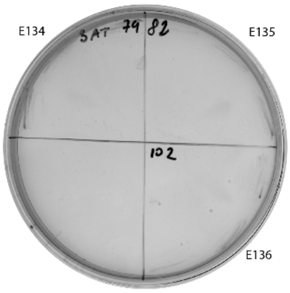

E137 AD-CG6792 BTB / BD-CG6792 BTB
E138 AD / BD-CG6792 BTB
E139 AD-CG32121 BTB / BD-CG6792 BTB
E140 AD-BCL6 BTB / BD-CG6792 BTB

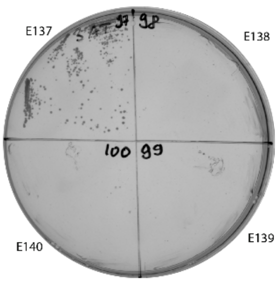

E141 BD-bab2 BTB / AD-KEN BTB
E142 BD-bab1 BTB / AD-KEN BTB
E143 BD-Batman BTB / AD-KEN BTB
E144 BD-CG34376 BTB / AD-KEN BTB

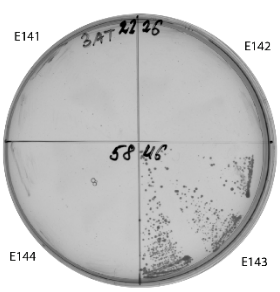

E145 AD-CG15725 BTB / BD-CG15725 BTB
E146 AD / BD-CG15725 BTB
E147 AD-CG32121 BTB / BD-CG15725 BTB
E148 AD-BCL6 BTB / BD-CG15725 BTB

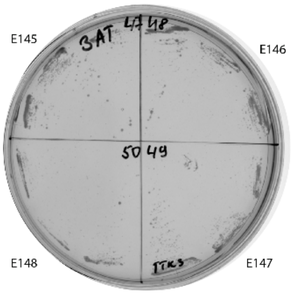

E150 BD-Mod(mdg4) BTB / AD-CG15725 BTB
E151 BD-Mod(mdg4) BTB / AD
E152 BD-Abrupt BTB / AD-CG15725 BTB
E153 BD-Abrupt BTB / AD

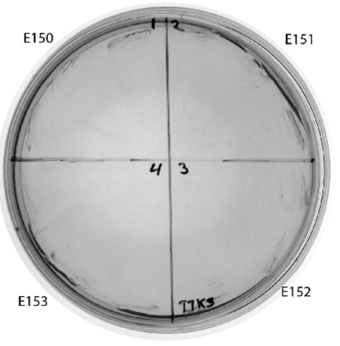

E154 BD-CG3726 BTB / AD-CG15725 BTB
E155 BD-CG3726 BTB / AD
E156 BD-CG12236 BTB / AD-CG15725 BTB
E157 BD-CG12236 BTB / AD

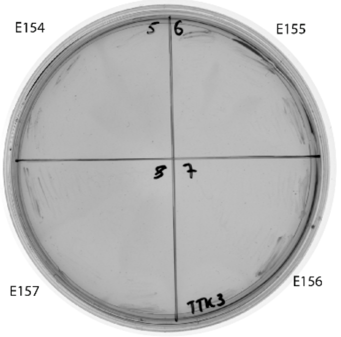

E158 BD-BTB-VII BTB / AD-CG15725 BTB
E159 BD-BTB-VII BTB / AD
E160 BD-Bab2 BTB / AD-CG15725 BTB
E161 BD-Bab2 BTB / AD

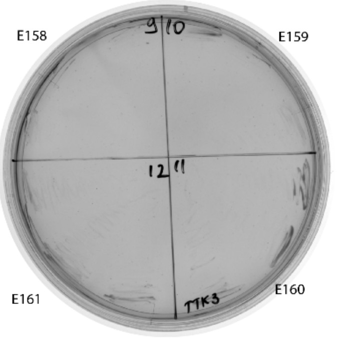

E162 BD-Bab1 BTB / AD-CG15725 BTB
E163 BD-Bab1 BTB / AD
E164 BD-Ribbon BTB / AD-CG15725 BTB
E165 BD-Ribbon BTB / AD

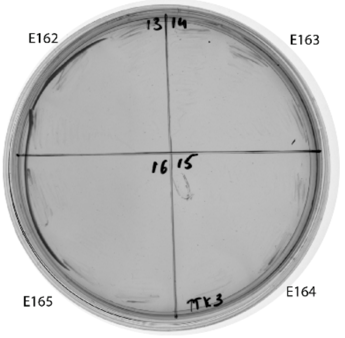

E166 BD-GAF BTB / AD-CG15725 BTB
E167 BD-GAF BTB / AD
E168 BD-TTK BTB / AD-CG15725 BTB
E169 BD-TTK BTB / AD

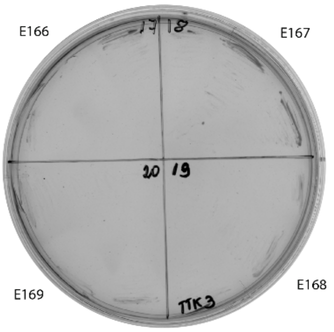

E170 BD-Pipsqueak BTB / AD-CG15725 BTB
E171 BD-Pipsqueak BTB / AD
E172 BD-Batman BTB / AD-CG15725 BTB
E173 BD-Batman BTB / AD

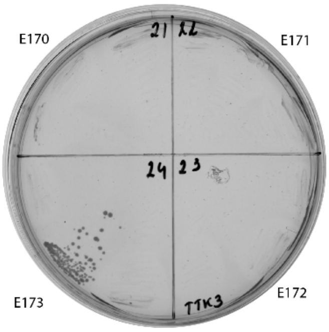

E174 BD-CG6118 BTB / AD-CG15725 BTB
E175 BD-CG6118 BTB / AD
E176 BD-CG15812 BTB / AD-CG15725 BTB
E177 BD-CG15812 BTB / AD

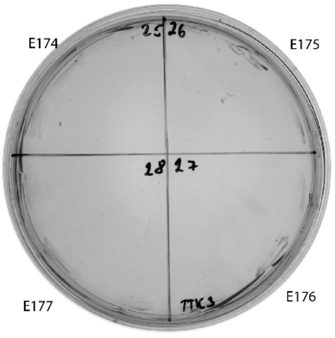

E178 BD-CG34376 BTB / AD-CG15725 BTB
E179 BD-CG34376 BTB / AD
E180 BD-Fruitless BTB / AD-CG15725 BTB
E181 BD-Fruitless BTB / AD

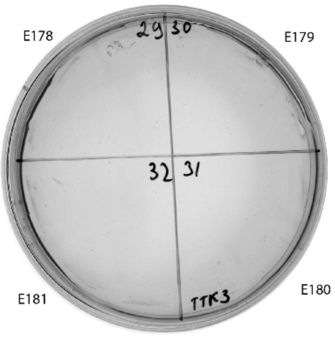

E182 BD-TKR BTB / AD-CG15725 BTB
E183 BD-TKR BTB / AD
E184 BD-CG8924 BTB / AD-CG15725 BTB
E185 BD-CG8924 BTB / AD

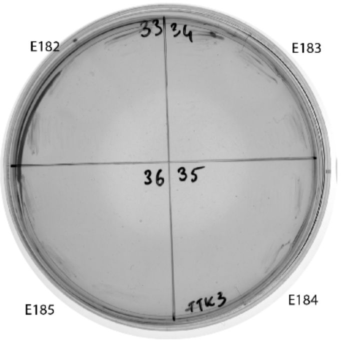

E186 BD-MAMO BTB / AD-CG15725 BTB
E187 BD-MAMO BTB / AD
E188 BD-BR-C BTB / AD-CG15725 BTB
E189 BD-BR-C BTB / AD

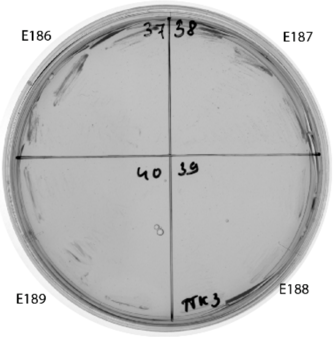

E190 BD-Chinmo BTB / AD-CG15725 BTB
E191 BD-Chinmo BTB / AD
E192 BD-CG6765 BTB / AD-CG15725 BTB
E193 BD-CG6765 BTB / AD

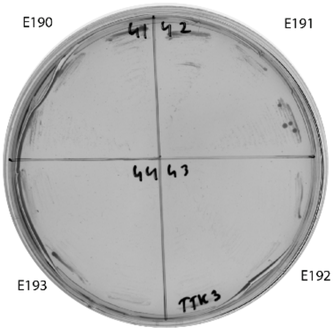

E194 BD / AD-CG15725 BTB
E195 BD / AD
E196 BD / AD-CG32121 BTB
E197 BD / AD-BCL6 BTB

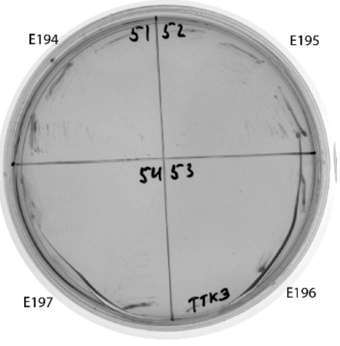

**Supplementary Figure S5. Superdex S200 size-exclusion chromatography of TTK-type BTB domains from *Drosophila*.** Positions of molecular weight markers (in kDa) are shown below. Molecular weights of monomers in kDa are shown in brackets.

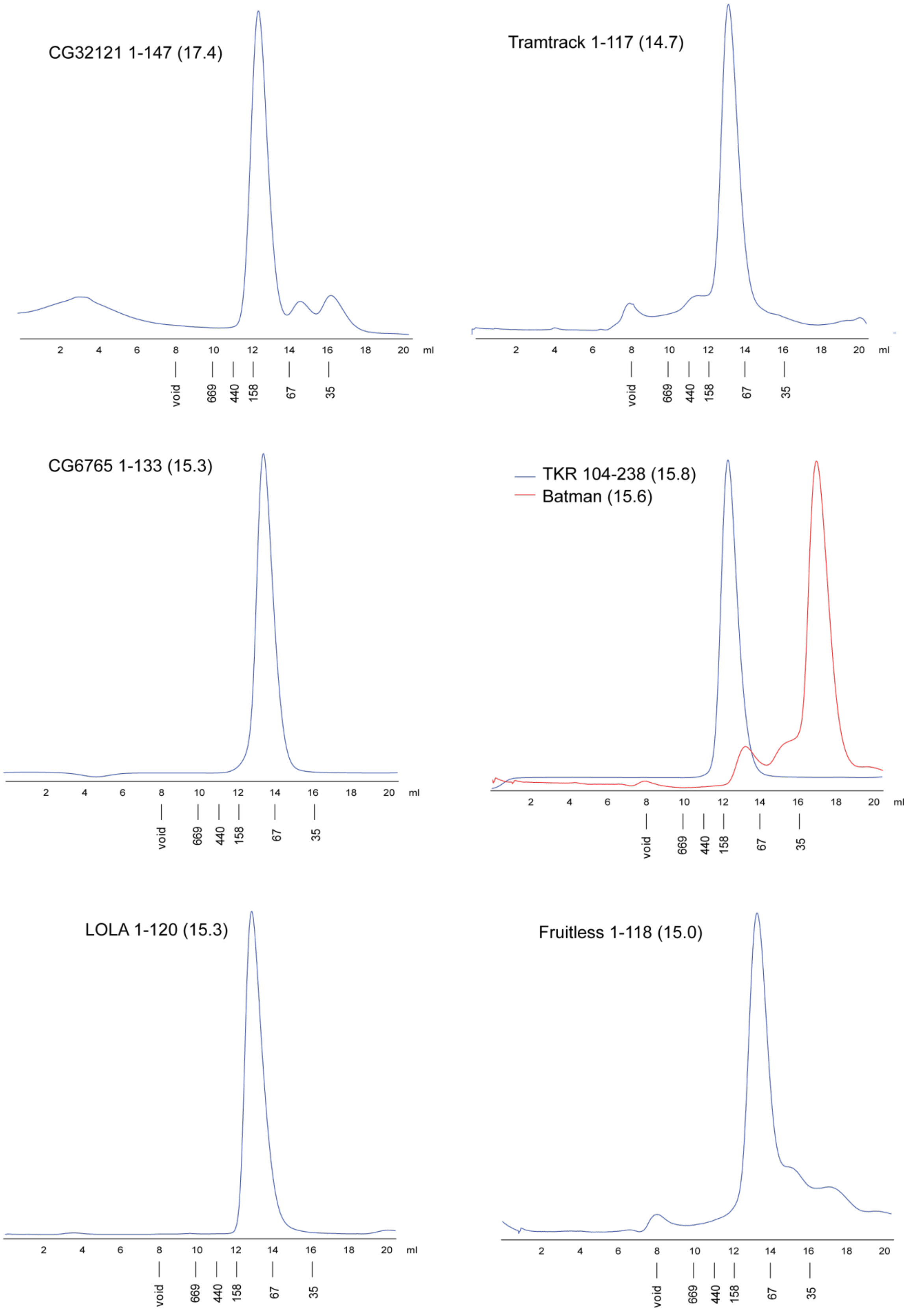

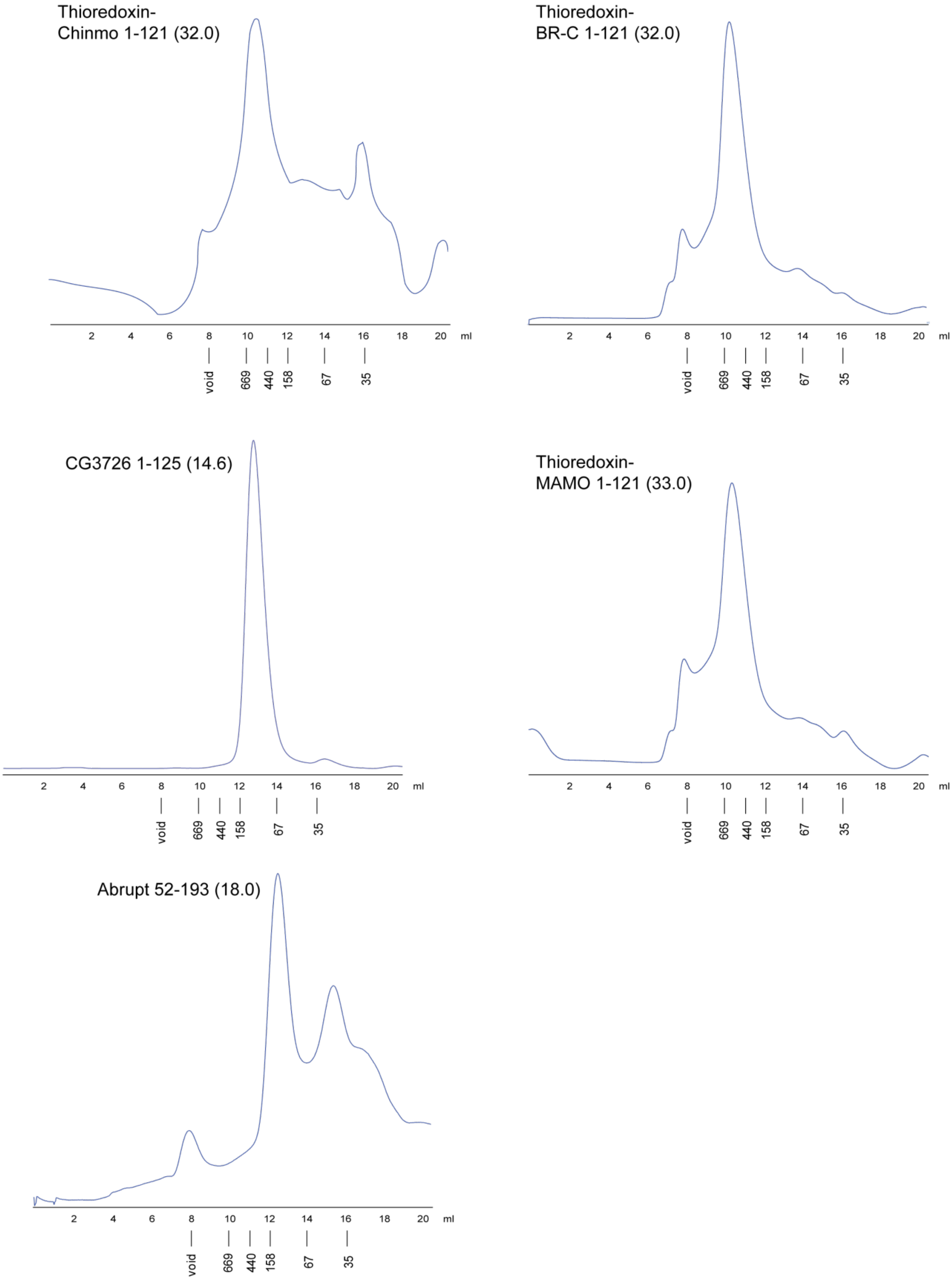
Thioredoxin-
Chinmo 1-121 (32.0)
Thioredoxin-
BR-C 1-121 (32.0)
CG3726 1-125 (14.6)
Thioredoxin-
MAMO 1-121 (33.0)
Abrupt 52-193 (18.0)
2 4 6 8 10 12 14 16 18 20 ml
void 669 440 158 67 35

**Supplementary Figure S6.** (A) Alphafold2 derived models of LOLA oligomers and their fits to the experimental SAXS data for the LOLA sample, shown along with the fit quality (chi2). **(B)** Alphafold2 derived models of CG6765 oligomers and their fits to the experimental SAXS data for the CG6765 sample, shown along with the fit quality (chi2). Note that the hexamers provided most satisfactory fits. All attempts to build 7-mers returned disconnected models, which were not considered in analysis.

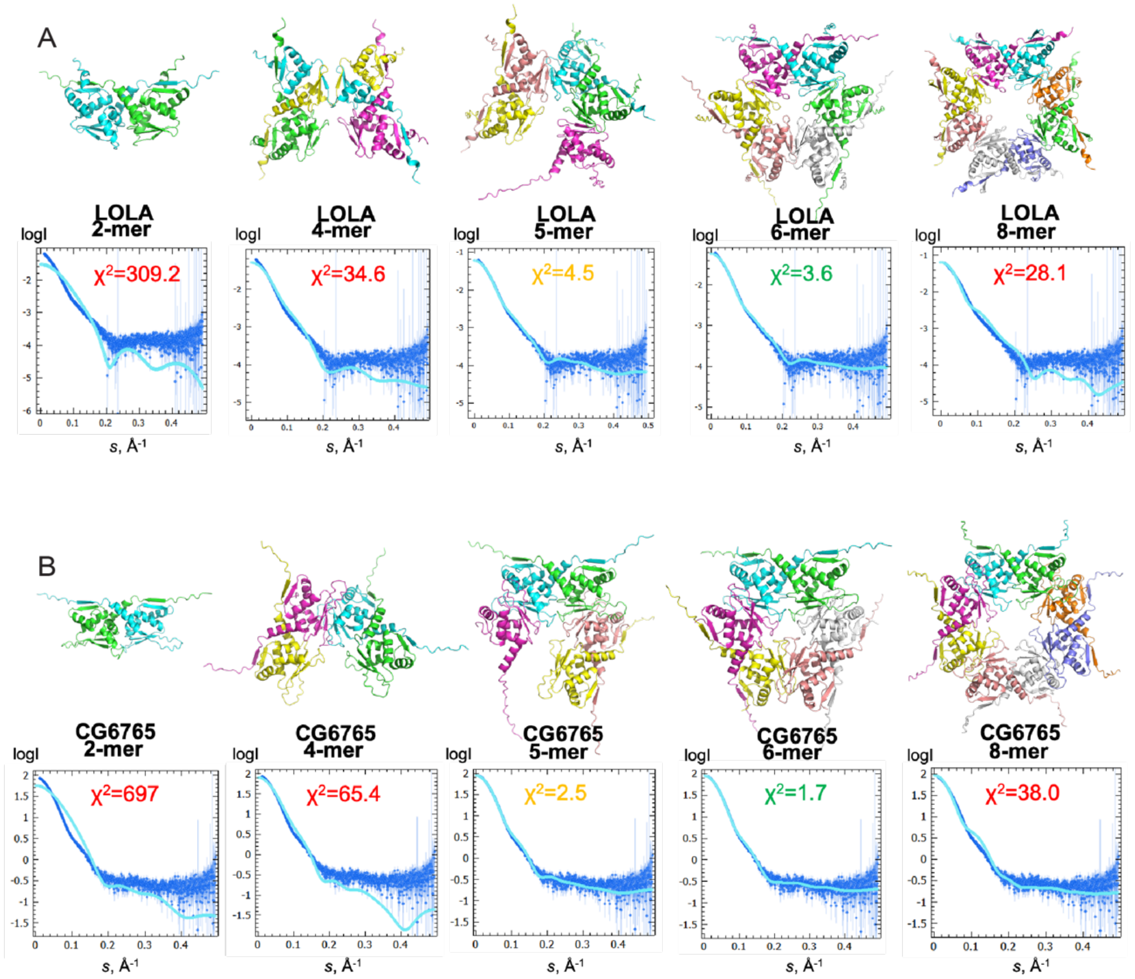

**Supplementary Figure S7. Molecular modeling of TTK-type BTB domains with AlphaFold. a** Models of hexameric assemblies of TTK-type BTB domains. Models are colored according to AlphaFold plDDT values. **b** Molecular modeling of Mod(mdg4) BTB domain hexamer with close-up view of dimer-dimer interaction interface. **c** Close-up view of dimer-dimer interaction interface of Alphafold models of LOLA, CG6765 and Batman hexamers.

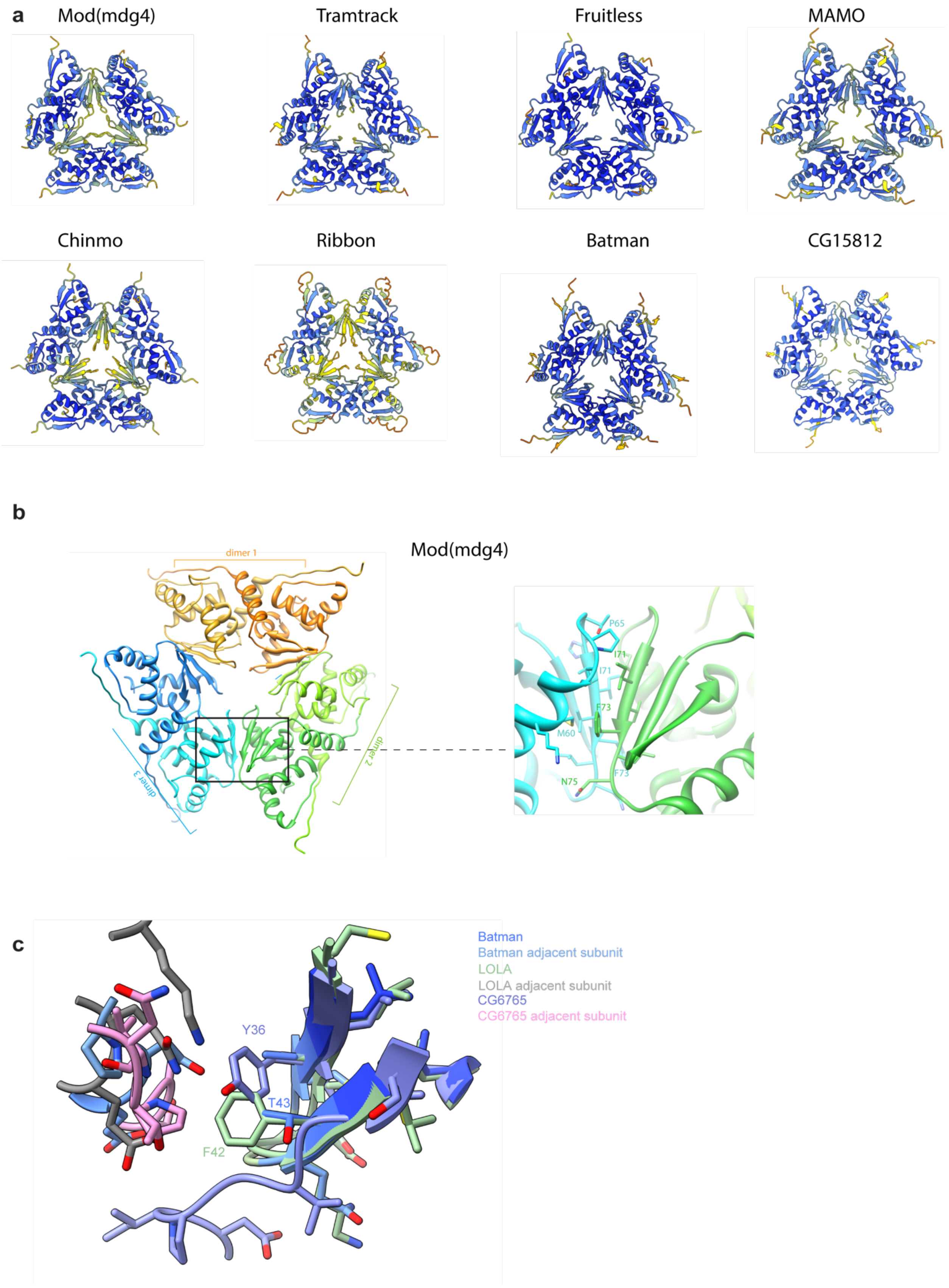

**Supplementary Figure S8. Testing the impact of single amino-acid substitutions in dimer-dimer interaction interface on the oligomerization status of TTK-type BTB domains.** Close-up views of AlphaFold-derived molecular models of dimer-dimer interaction interfaces and effect of single amino-acid substitutions on the oligomerization status of BTB domains of Mod(mdg4) **(a)**, LOLA **(b)**, and CG6765 **(c)** studied with Superdex S200 size-exclusion chromatography. Amino acids subjected to mutagenesis are shown in red. Positions of molecular weight markers are shown below. Molecular weights of monomers are shown in brackets. Further details are shown in Supplementary Figures S9, S10, and S11. **(d)** The self-interaction ability studied with a yeast two-hybrid assay. Growth assay plates are shown in Supplementary Figure S12. asterisk indicates that the interaction was observed only in case when wild-type BTB was BD-fused. Double asterisk indicates that result is unreliable due to high self-activatory activity.

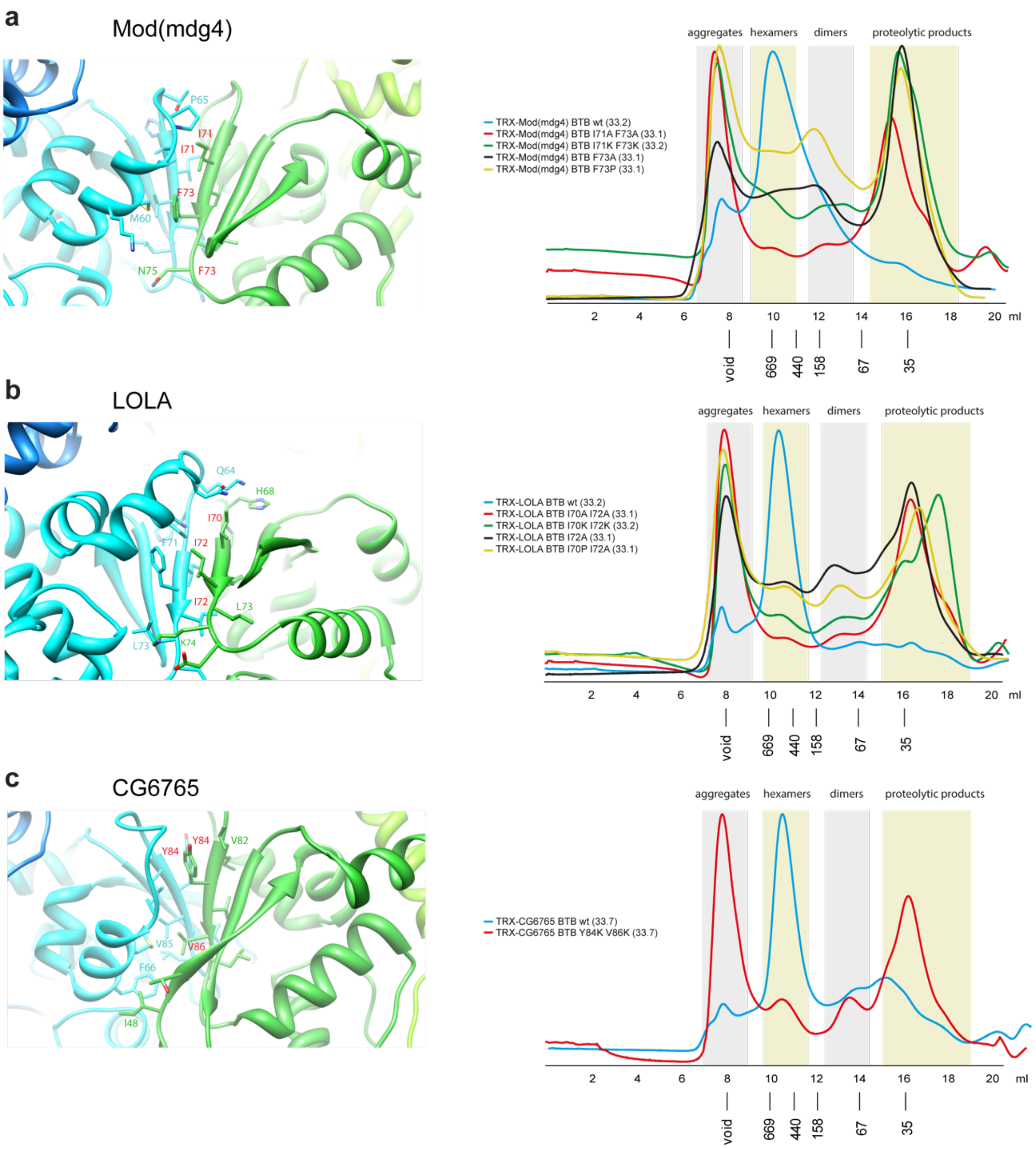

**d**

| | $LOLA^{I70A;I72A}$ | $LOLA^{I70K;I72K}$ | $LOLA^{I70A}$ | $MOD^{I71A;F73A}$ | $MOD^{I71K;F73K}$ | $MOD^{F73P}$ |
|---|---|---|---|---|---|---|
| Self-interaction control | + | –** | + | + | – | + |
| Interaction with BTB wt | +/-* | – | + | + | + | + |

**Supplementary Figure S9. Superdex S200 size-exclusion chromatography of Thioredoxin-tagged LOLA BTB domains bearing mutations in dimer-dimer interaction interface.** Positions of molecular weight markers (in kDa) are shown below. Molecular weights of monomers in kDa are shown in brackets. The gel showing the content of SEC fractions peaks of alanine mutant is shown below, molecular weight markers are shown at the left.

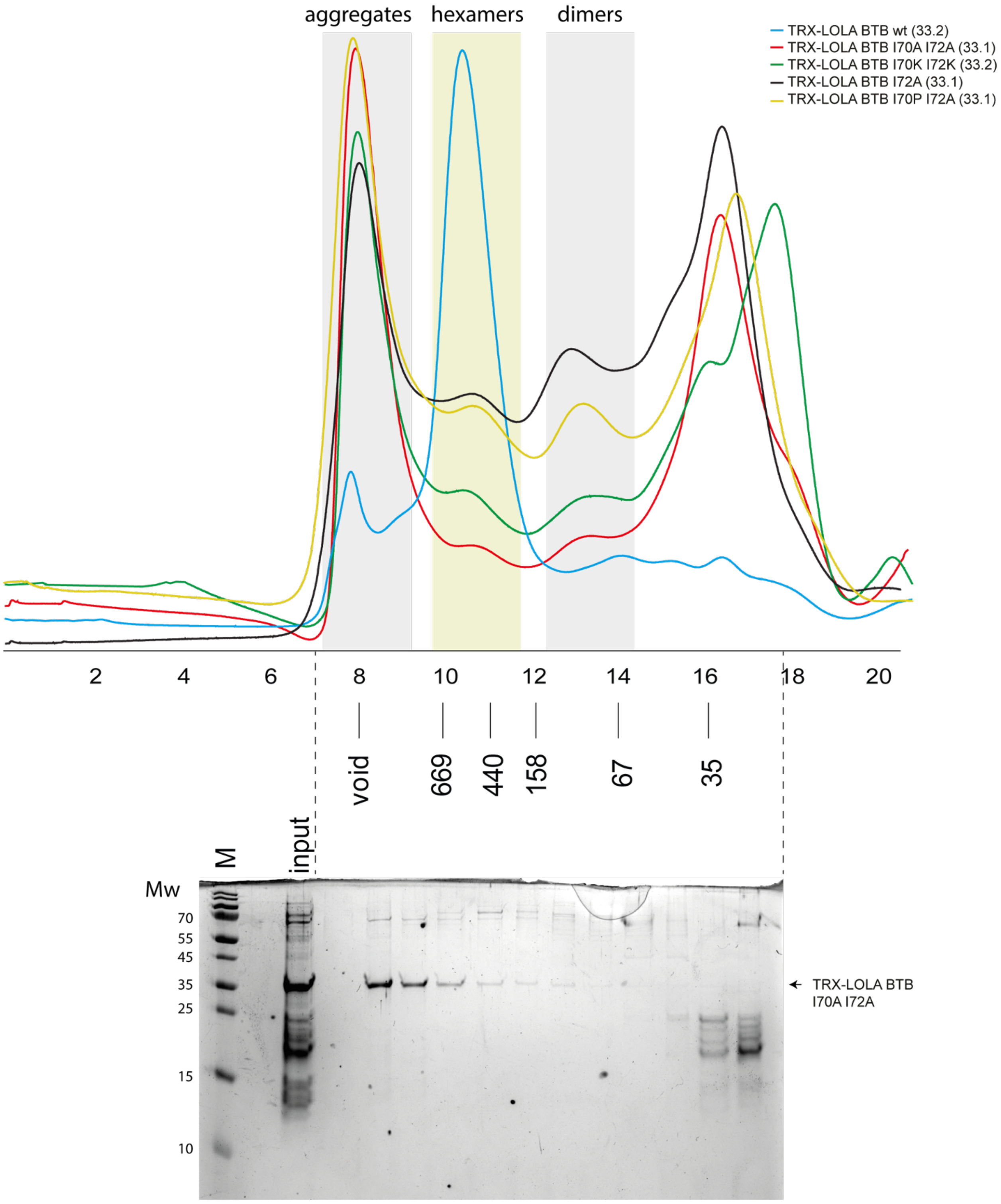

**Supplementary Figure S10. Superdex S200 size-exclusion chromatography of Thioredoxin-tagged Mod(mdg4) BTB domains bearing mutations in dimer-dimer interaction interface.** Positions of molecular weight markers (in kDa) are shown below. Molecular weights of monomers in kDa are shown in brackets. The gel showing the content of SEC fractions peaks of alanine mutant is shown below, molecular weight markers are shown at the left.

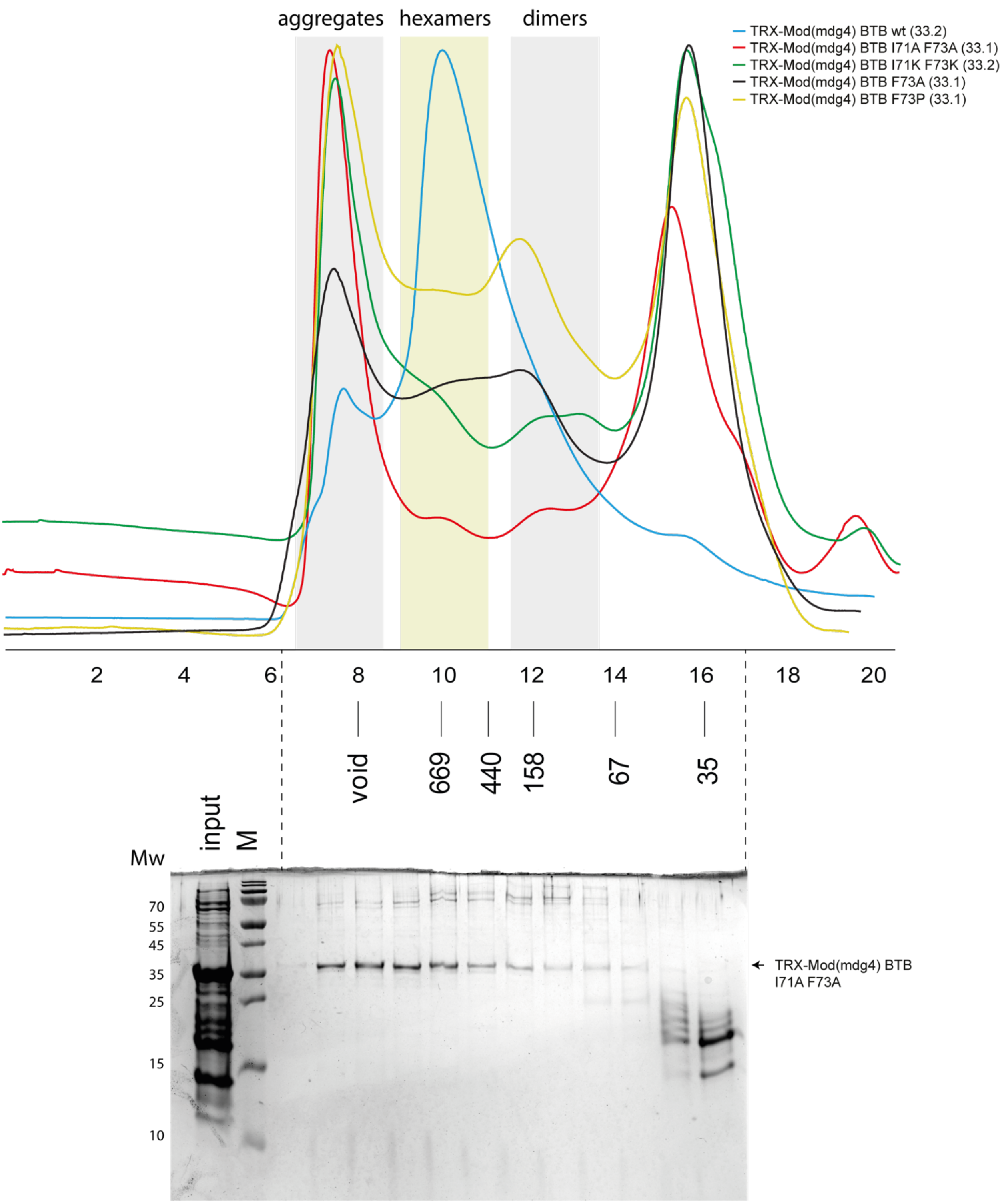

**Supplementary Figure S11. Superdex S200 size-exclusion chromatography of Thioredoxin-tagged CG6765 BTB domains bearing mutations in dimer-dimer interaction interface.** Positions of molecular weight markers (in kDa) are shown below. Molecular weights of monomers in kDa are shown in brackets. The gel showing the content of SEC fractions peaks of mutant protein is shown below, molecular weight markers are shown at the left.

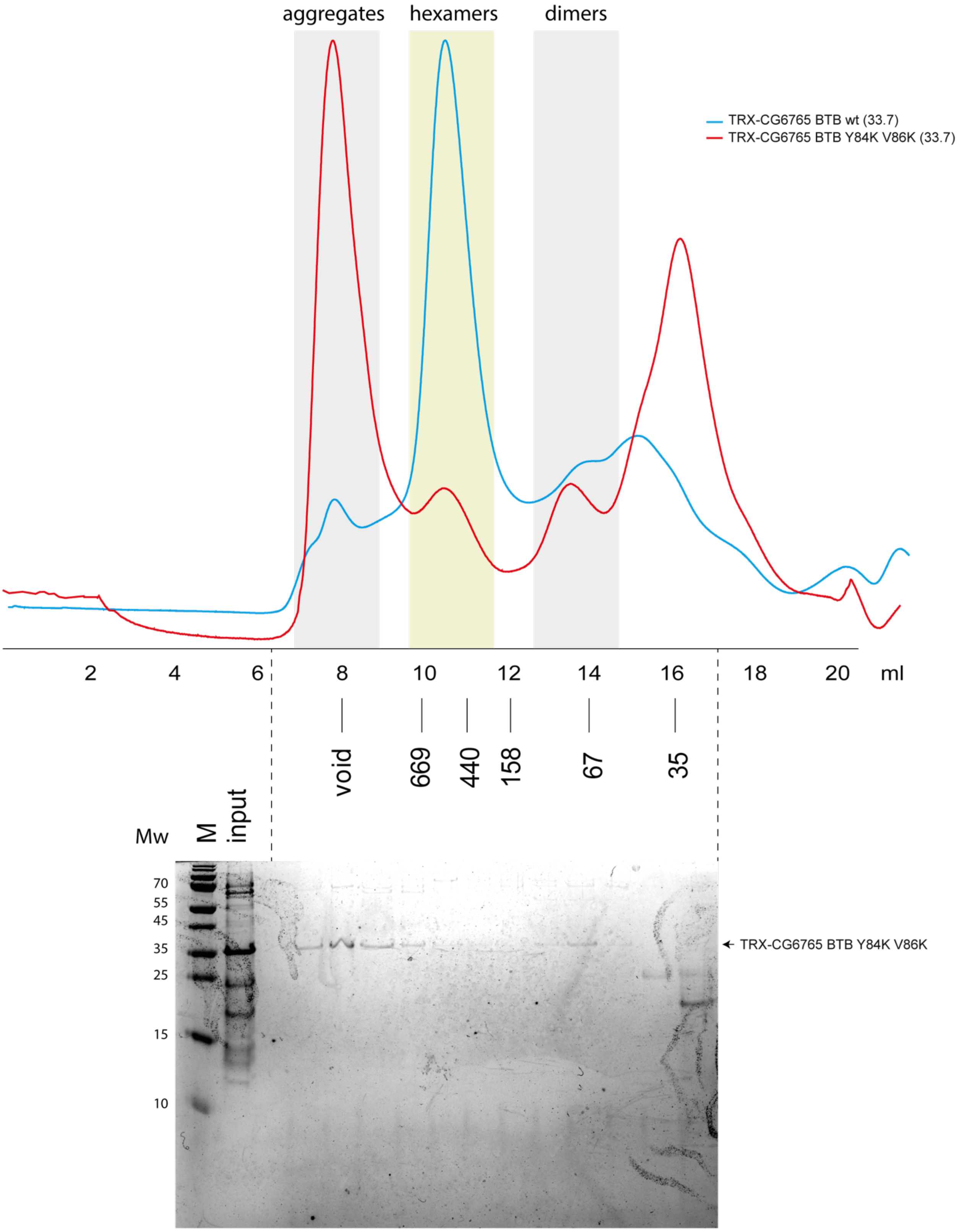

**Supplementary Figure S12. Results of yeast two-hybrid assays of the impact of point mutations at the dimer-dimer interaction interface.** Growth assay plates without histidine are shown (yeasts are unable to grow on this medium in the absence of interaction). AD stands for Activation Domain, BD – for DNA-Binding Domain of GAL4 protein.

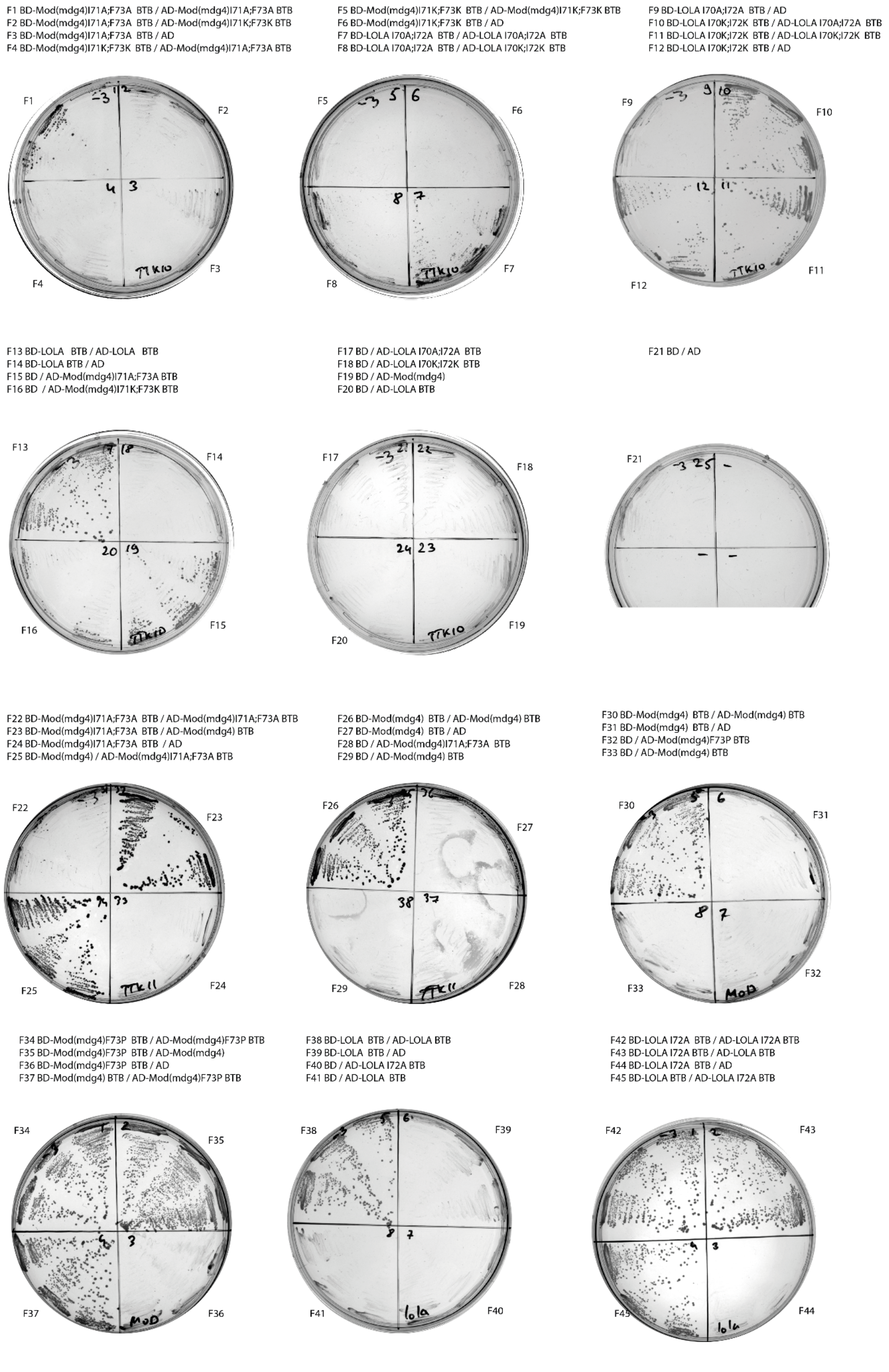
